# CRISPR inhibition of activity-dependent Arc expression in the adult mouse brain has limited effects on plasticity in visual cortex and nucleus accumbens

**DOI:** 10.64898/2026.04.26.720914

**Authors:** Aaron T. Halvorsen, Arthy Narayanan, Grace M. Link, Thomas J. Wagner, Ashleigh K. Waterman, Mara H. Cowen, Katherine R. Tonn Eisinger, Lindsey L. Glickfeld, Anne E. West

## Abstract

Neuronal activity drives long-lasting change in circuit function by inducing the expression of gene products that modulate the function of synapses. *Arc* is one of the most robustly activity-regulated neuronal genes, being induced broadly in neurons by a wide range of physiologically-relevant stimuli. The ability of Arc to promote internalization of AMPA-type glutamate receptors plays an important role in activity-dependent refinement of synaptic connectivity during development. However, the consequences of Arc induction for circuit plasticity in the adult brain are less well understood. We reasoned that we could test the requirement for experience-induced Arc expression in downstream plasticities by disrupting regulatory elements that mediate the inducibility of *Arc* transcription. To achieve this goal, we developed and validated a CRISPR-based inhibition strategy to conditionally block stimulus-induced expression of Arc in specific regions of the brains of male and female adult mice in vivo. We show that recruiting transgenic dCas9-KRAB to decrease transcriptional activity of either the promoter or the synaptic activity-regulated enhancer of the *Arc* gene is sufficient to block Arc protein expression in visual cortex (V1) following light exposure or in the nucleus accumbens (NAc) following cocaine administration. However, loss of Arc in V1 failed to alter plasticity of orientation selectivity and loss of Arc in NAc did not block cocaine conditioned place preference or novel object recognition memory. These data show that the relationship between Arc induction and plasticity is not universal and suggest that additional contextual factors determine the functional consequences of Arc induction for plasticity.

**Significance Statement:** Arc is a neuronal activity-regulated gene whose expression is robustly induced in the adult brain by a wide range of stimuli that drive plasticity. However, the functional requirements of Arc induction for brain plasticity are incompletely understood. Here, we show that we can use transgenic mice expressing the CRISPR-based transcriptional repressor dCas9-KRAB to block stimulus-induced expression of Arc in specific regions of the adult mouse brain. Despite highly effective inhibition of Arc induction, we found that many forms of plasticity remained intact. These data deepen understanding of the contextual importance of activity-induced Arc expression in the adult brain.

## Introduction

*Arc* (activity-regulated cytoskeleton-associated protein) transcription is rapidly and transiently induced in both developing and adult neurons by neuronal activity (Lyford et al., 1995). The observation that both *Arc* mRNA and Arc protein are trafficked to active synapses in hippocampal dendrites following stimuli that induce long-term potentiation (LTP) first suggested the possibility that Arc induction might directly couple activity-dependent transcription with synapse plasticity (Steward et al., 1998). This model of experience-induced Arc as a nucleus-to-synaptic mediator of memory was supported by germline knockout (KO) of Arc, which disrupted long-term, but not short-term, memory in behavioral tasks (Plath et al., 2006).

Though Arc was first anticipated to mediate LTP as a mechanism for its role in memory, subsequent mechanistic studies showed that Arc promotes the endocytosis of AMPA-type glutamate receptors (AMPARs) especially at weak synapses, and that loss of Arc impairs hippocampal long-term depression (LTD) rather than LTP (Chowdhury et al., 2006; Kyrke-Smith et al., 2021; Plath et al., 2006; Shepherd et al., 2006). This cellular function of Arc in AMPAR endocytosis is required for experience-dependent refinement of synaptic connectivity during critical periods of sensory-dependent brain development (Jenks et al., 2017; McCurry et al., 2010; Mikuni et al., 2013). In the adult brain, regional KO or knockdown (KD) of Arc after the period of developmental wiring recapitulates certain features of developmental Arc loss, though the mechanisms of Arc action in these studies are less clear. For example, both developmental and adult depletion of Arc impair hippocampal memory in mice (Coda et al., 2025; Gao et al., 2018), however changes in hippocampal oscillations that accompany learning are only impaired with developmental Arc loss. Depletion of Arc in the developing or adult mouse visual cortex increases the number of binocular neurons (Jenks & Shepherd, 2020), though the relationship to experience-dependent synapse plasticity is not known.

We reasoned that if the experience-dependent regulation of Arc is broadly required for plasticity in the adult brain, then selectively disrupting the inducibility of *Arc* transcription should facilitate study of this mechanism. Neuronal experience-inducible genes are targets of the stimulus-activated transcription factors (TFs) CREB, MEF2, and SRF (West, 2025). When these TFs are phosphorylated by activity-regulated kinases, they increase the likelihood that RNA polymerase II is recruited to the promoters of their target genes.

Interestingly, these activity-regulated TFs bind the *Arc* gene at a single synaptic-activity regulated enhancer element ∼7kb upstream of the transcriptional start site rather than the *Arc* promoter (Kawashima et al., 2009; Waltereit et al., 2001). We hypothesized that we could use this spatial separation of regulatory elements to control the basal versus the activity-inducible transcription of *Arc* by targeting CRISPR inhibition (CRISPRi) machinery to either the promoter or enhancer of *Arc* respectively. CRISPRi uses an enzymatically dead version of the bacterial Cas9 protein (dCas9) as an RNA-guided, DNA binding tether to anchor the repressor domain of the TF KRAB (dCas9-KRAB) to any given gene regulatory element (Gemberling et al., 2021), thus offering an efficient strategy to conduct functional genomics (Bach et al., 2024; Joo et al., 2016).

Here, we design CRISPRi guide RNAs (gRNAs) to the *Arc* promoter or enhancer and demonstrate we can robustly block activity-inducible *Arc* RNA and Arc protein expression both in primary neuronal culture and in the adult brain of conditional dCas9-KRAB transgenic mice. We disrupt Arc expression in the visual cortex (V1) or nucleus accumbens (NAc) of adult mice to test the requirement for Arc regulation in two different cell types and circuit plasticity contexts: (1) representational drift of visual orientation selectivity in glutamatergic neurons of V1 and (2) cocaine-induced neural ensemble activity of medium spiny neurons in the NAc. Surprisingly, we found minimal functional consequences of adult Arc loss in either setting. These findings challenge the model of Arc as a universal mediator of activity-induced synapse plasticity and suggest that additional cellular mechanisms shape the cell type- or synapse-specific consequences of Arc induction in the adult brain.

## Materials and Methods

### Mice

All experiments were performed in accordance with an approved protocol from the Institutional Animal Care and Use Committee (IACUC) at Duke University. Mice were housed under standard, temperature-controlled housing conditions with ad libitum food and water and male and female mice were used for all experiments. Rosa26-LSL-dCas9-KRAB (dCas9-KRAB, strain #033066, RRID:IMSR_JAX:033066) and Rosa26-LSL-dCas9-p300 mice (dCas9-p300, strain #033065, RRID:IMSR_JAX:033065) were used to modulate Arc expression (Gemberling et al., 2021). Mice homozygous for the transgenes were crossed with C57BL/6J mice (Jackson Labs strain #000664, RRID:IMSR_JAX:000664) to produce heterozygous male and female mice, which were used for all in vivo experiments. Timed-pregnant CD1-IGS mice (strain #022; RRID: IMSR_CRL022) were purchased from Charles River Labs for embryonic culture experiments. For Arc induction in visual cortex, mice were dark-housed for 4 days prior to light exposure for the times described. Behavioral testing was performed during the light portion of a 14:10 light/dark cycle housing by an experimenter blind to genotype.

### Embryonic neuronal cell culture

Timed-pregnant CD1-IGS mice (Charles River strain #022; RRID: IMSR_CRL022) were anesthetized with isoflurane and rapidly decapitated. Brains from male and female E16.5 embryos were rapidly removed, the cortices were dissected and enzymatically dissociated in papain (Worthington, Cat# LS003126) for 10 min, and the cells were mechanically dissociated and initially plated in DMEM with 10% added calf serum. For the immunocytochemistry experiments, the neurons were plated on poly-D-lysine (PDL) and laminin coated coverslips at a density of 125-250K/coverslip. For the RT-qPCR experiments, the neurons were plated on PDL coated 24 well tissue culture dishes at a density of 500K/well. Four hours after plating, media was switched to Neurobasal plus B27 (Gibco, Cat# 21103-049, Cat# 17504-044). On day in vitro 1 (DIV1), conditioned media was removed, and the neurons were infected with lentivirus at a multiplicity of infection (MOI) of 1 in plain Basal Media Eagle (Sigma Aldrich) with 0.4mg/ml polybrene for 6 hr, after which the virus was removed, and the conditioned media was replaced. For the induction of Arc, on DIV7, the neurons were stimulated with 50ng/mL BDNF (Peprotech, Cat# 450-02) for the periods of time noted in the text prior to harvesting.

### Viruses

Lentiviral plasmids were generated by first restriction digesting and removing the GFP from pLV hU6-gRNA(anti-sense) hUbC-VP64-dCas9-VP64-T2A-GFP (Addgene #66707) and pLV hU6-sgRNA hUbC-dCas9-KRAB-T2A-GFP (Addgene #71237) using KpnI and cloning in an mCherry fluorophore at that site. The gRNA sequences (**Table 1**) were cloned into the BsmBI restriction site. These constructs were packaged as lentiviruses using HEK293T cells with pCMV-VSVG (Addgene #8454) and Δ8.91 (Addgene #187441) plasmids following standard procedures, purified by ultracentrifugation, and functionally titered in both HEK293T cells and cultured cortical neurons using imaging for mCherry. Adeno-associated viruses used for in vivo experiments were produced by the Duke University Viral Vector Core facility and included pGP-AAV-syn-jGCaMP7f-WPRE (Addgene #104488) and AAV:ITR-U6-sgRNA(backbone)-hSyn-Cre-2A-EGFP-KASH-WPRE-shortPA-ITR (Addgene #60231). An additional dsRed virus was made by replacing the GFP-WPRE of Addgene #60231 with dsRed. All AAV viruses had AAV1 serotype. Viruses with gRNAs were targeted to either *LacZ*, the *Arc* promoter, or the *Arc* enhancer. Two gRNA viruses were pooled together in a 1:1 ratio for the promoter and the enhancer (see **Table 1** for sequences and details). Viral titers of AAVs averaged around 3E13 vg/mL for each virus.

**Table 1.**
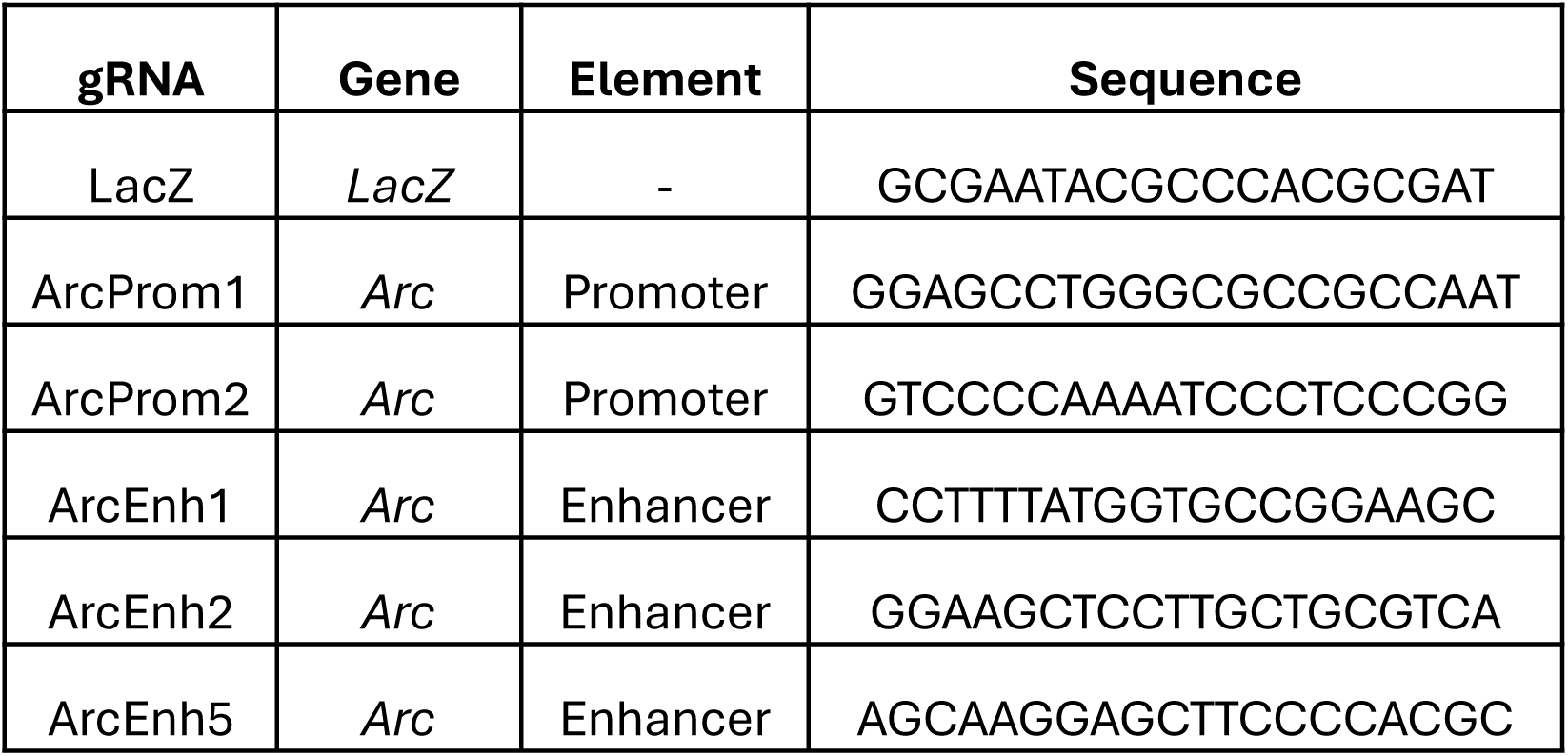
gRNA sequences used in viruses. Two viruses were pooled for promoter and enhancer targeting. Promoter gRNAs were equivalent in vitro and in vivo. For the enhancer, ArcEnh1 + Arc Enh2 were used in vitro, and ArcEnh1+ ArcEnh5 were used in vivo.

### RNA Extraction and quantitative RT-PCR

RNA was harvested with TRIzol (Invitrogen) and cDNA was reverse transcribed by random hexamer priming with Superscript II (Invitrogen, 18064014). Quantitative SYBR green PCR was performed on a Thermo-Fisher/Applied BioSystems QuantStudio3 quantitative real-time PCR system using the primers in **Table 2**.

**Table 2.**
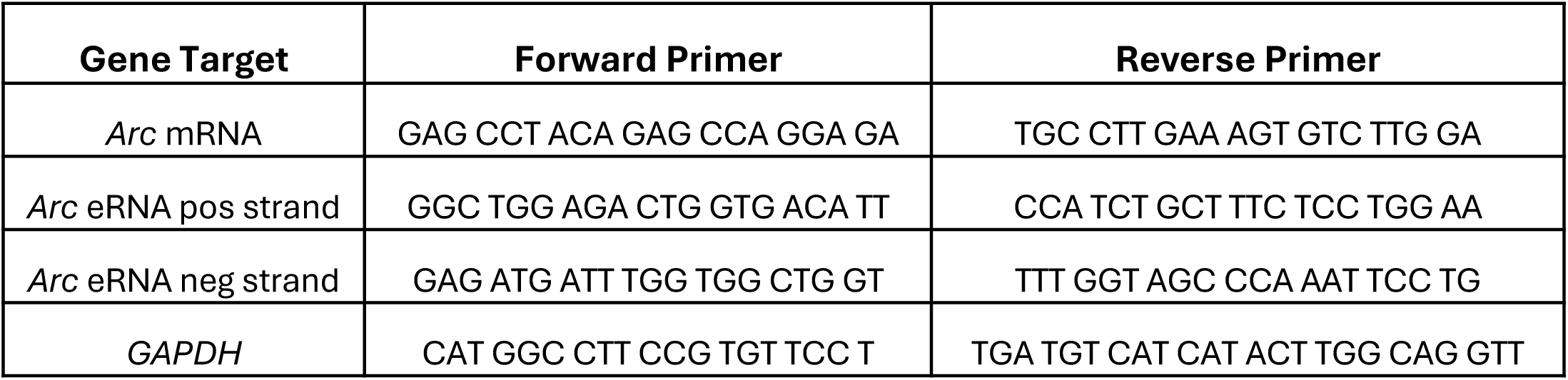
Primers used in qPCR experiments.

### Immunocytochemistry

Coverslips were fixed with 4% paraformaldehyde (PFA) and 4% sucrose for 10 min and blocked and permeabilized for 1 hr in PBS with 1% Tx-100 and 10% goat serum prior to overnight (O/N) incubation at 4°C in PBS with 0.3% Tx-100, 10% goat serum, and rabbit anti-Arc antibody (1:1000, Synaptic Systems 156003). Cells were washed 3X with PBS and incubated with goat anti-rabbit Cy 2 (1:500, Jackson Immuno, 305-227-003) for 3 hr at room temperature (RT) prior to rising with PBS, labeling nuclei with Hoechst (1:1000 Thermo Fisher Scientific, H1399) and mounting in Vectashield (VectorLabs H1000-10).

### Western blotting

2x Laemmli Sample Buffer (Bio-Rad, Cat# 1610737) and 2-mercaptoethanol were added to each sample and boiled for 10 min prior to separation by SDS-PAGE on a 12% gel and transfer to nitrocellulose. Blots were blocked in 3% bovine serum albumin and 0.2% sodium azide in 1x TBST for 1 hr at RT. Rabbit anti-Arc (1:1000, Synaptic Systems 156003) and mouse anti-actin (1:7000, Sigma-Aldrich MAB1501) were added and blots shaken in solution antibody solution at 4°C O/N. Blots were washed 3 times with TBST then incubated with same blocking solution with secondary antibodies goat anti-rabbit (1:5000, Biotium Cat# 20067), and goat anti-mouse (1:5000, Biotium Cat# 20065) for 1 hr at RT. The blots were washed in TBST before drying and imaging on a LiCor Odyssey.

### Stereotactic surgery

Viral surgeries were performed on male and female mice at age 10-18 weeks. Nucleus accumbens (NAc) and visual cortex (V1) injections consisted of a bilateral injection of 0.5µl virus (NAc) or 0.25µl (V1). For within-mouse control experiments, mice were injected with a *LacZ* control gRNA virus into the left hemisphere, and either *Arc* promoter or *Arc* enhancer gRNA viruses were injected into the right hemisphere. Mice were anesthetized using isoflurane and given a 5mg/kg injection of Meloxicam. Small burr holes were made in the skull with a drill (Dremel). For the NAc injections, 0.5ul of virus was delivered via Hamilton syringe at the following coordinates relative to Bregma: AP +1.6mm, ML +/- 0.7mm, and DV -4.5mm from the dura. Virus was injected manually at a rate of 0.1µl per minute. For the V1 injections, 0.1ul of virus was delivered at +/- 2.55mm lateral from lambda, directly anterior to the lambdoid suture, and ∼250µm below the pia. Virus was injected at 30µl/min through an UltraMicroPump (World Precisions Instruments). All experiments occurred >2 weeks after surgery to allow for the mouse to recover and for the virus to express.

### Immunohistochemistry

Mice were anesthetized with isoflurane, then transcardially perfused with PBS followed by 4% PFA in PBS. Brains were collected and post-fixed O/N in 4% PFA before being sunk O/N in 30% sucrose in PBS. Brains were sectioned coronally on a freezing microtome into 40µm slices and stored in PBS at 4°C. Slices were blocked and permeabilized in PBS with 1% Triton X-100 and 10% NGS for 1 hr, then incubated as floating sections O/N at 4°C in PBS with 0.3% Triton X-100, and 10% Normal Goat Serum with antibodies, including rabbit anti-Arc (1:1000, Synaptic Systems 156003), rabbit anti-Fos (1:1000, Synaptic Systems 226008), mouse anti-NeuN (1:200, EnCor MCA-1B7), and rabbit anti-H3K27ac (1:1000, abcam 177178). After washing with PBS, slices were incubated with appropriate secondary antibodies including goat anti-rabbit Cy2 (1:500, Jackson Immuno, 305-227-003) goat anti-rabbit Cy3 (1:500, Jackson Immuno, 111-165-144), goat anti-mouse Cy3 (1:500, Jackson Immuno, 115-165-003), goat anti-rabbit 594 (1:500, ThermoFisher, #A-11012), goat anti-rabbit Cy5 (1:500, Jackson Immuno, 111-175-144) for 1 hr at RT. Slices were washed 3x in PBS, nuclei labeled with Hoechst (1:1000 Thermo Fisher Scientific, H1399), and coverslipped and mounted in Vectashield (VectorLabs H1000-10).

### Immunohistochemistry quantification

Z-stack images were taken on a Leica SP8 confocal microscope. Image settings were kept constant within each experiment. Images consisted of 3 channels: Hoechst (nuclear stain), GFP (from virus), and Arc or Fos. All channels were max projected in FIJI/ImageJ to prepare for processing.

For the visual cortex images, the machine learning software Ilastik (Berg et al, 2019) was trained on max projected GFP channels for sample slices representing all conditions to identify cells. Training was done by hand by assigning example pixels as either GFP+ or GFP-pixels. Following training, all images were processed through the algorithm and exported as “Simple Segmentation” tif files, which were converted to masks in FIJI and processed using the Watershed function to separate any overlapping nuclei. The Analyze Particles function of FIJI was used to count cells, filtering by size and circularity. The resulting ROIs were overlaid on the Arc channel for each slice, and the cell number and the cell integrated densities of Arc immunofluorescence were measured for each cell. All layers were analyzed in the promoter mice, and all layers except layer 6 were analyzed in the enhancer mice. For the NAc, *Arc* enhancer images Ilastik was trained to look for Arc+ or Fos+ cells within the region of GFP expression. For the NAc *Arc* promoter images, the Arc+ and Fos+ cells were counted by hand by an experimenter blind to treatment. In both cases, the counts were normalized by the area of the GFP infected region.

### Headpost implantation surgery

Calcium imaging experiments were conducted at 16-39 weeks of age. Headpost and cranial window implantation were performed no earlier than 11 weeks. Animals were implanted with a titanium headpost and 5 mm cranial window as previously described (Goldey et al., 2014). Briefly, the headpost was secured using Metabond (Parkell) and a 5 mm craniotomy was made over the left hemisphere (center: 2.8 mm lateral, 0.5 mm anterior to lambda) allowing implantation of a glass window.

### Retinotopic mapping

Following at least 7 d recovery from the headpost implantation surgery, mice were gradually habituated to head restraint. After habituation, mice underwent retinotopic mapping using intrinsic autofluorescence imaging to locate V1 for viral injection. The brain was illuminated with white light (Lumen Dynamics, X-Cite 120) with a 472 ± 30 nm band pass filter (Edmund Optics), and emitted light was measured through a green and red filter (500 nm longpass). Drifting gratings were presented on a monitor positioned at 45° relative to the body axis, and stimuli were shown at 3 positions (Elevation: -10 deg, Azimuth: -30, 0, and 30 deg, 45° diameter with a gaussian mask, drifting at 2 Hz, 10 s duration, 10 s inter-trial interval (ITI)) to activate locations in the contralateral visual field. Images were collected using a CCD camera (Rolera EMC-2, QImaging) at 2 Hz through a 5x air immersion objective (0.14 numerical aperture (NA), Mitutoyo), using Micromanager acquisition software (NIH). Images were analyzed in ImageJ (NIH) to measure changes in fluorescence (dF/F; with F being the average of all frames). Injections were targeted to the region of V1 driven by the center position.

### Viral injections for two-photon imaging

The mice used for two-photon imaging underwent additional surgery for viral injection. After anesthesia with isoflurane, the cranial window was removed. AAV1.Syn.GCaMP7f.WPRE mixed with AAV encoding *LacZ*, *Arc* Enhancer, or *Arc* Promoter gRNA in a 1:1 ratio was injected via a glass micropipette mounted on a Hamilton syringe. One hundred nanoliters of virus were injected at ∼250 µM below the pia (30 nL/min); the pipette was left in the brain for an additional 3 min to allow the virus to infuse into the tissue. Following injection, a new coverslip was sealed in place with Metabond. We then waited a minimum of one week for viral expression to mature before performing two-photon experiments.

### Two-photon imaging

Images were collected using a two-photon microscope controlled by Scanbox software (Neurolabware). A Mai Tai eHP DeepSee laser (Newport) was directed into a modulator (Conoptics) and raster scanned on the visual cortex using resonant galvanometers (8 kHz; Cambridge Technology) through a 16X (0.8 NA, Nikon) water-immersion lens at a frame rate of 15 Hz. Emitted photons were directed through a green (510 ± 42 nm band filter; Semrock) or red filter (607 ± 70 nm band filter; Semrock) onto GaAsP photomultipliers (H10770B-40, Hamamatsu). At the start of each experiment, we used an excitation wavelength of 1040 nm to visualize dsRed fluorescence, allowing identification of red gRNA expressing cells. All functional imaging used an excitation wavelength of 920 nm. Data were collected at 140 – 250 µM below the cortical surface. During imaging experiments, mice were head-fixed and allowed to freely run on a cylindrical treadmill. Running speed was monitored with a digital encoder (US Digital). For each mouse we performed three sequential imaging sessions (D1, D3 and D7). In a subset of mice (10/14), a second imaging plane was collected for an additional three imaging sessions.

### Visual stimulus presentation

Visual stimuli were presented on a 144-Hz (Asus). The monitor was calibrated with an i1 Display Pro (X-rite) for mean luminance at 50 cd/m2 and positioned 21 cm from the eye. Stimuli were generated and displayed using MWorks (The MWorks Project). At the beginning of each session, we performed a retinotopy (9 positions, 30 deg diameter gabor grating, 15 deg spacing in azimuth and elevation) to position the monitor such that the receptive fields of the imaged neurons were centered on the screen. Then, full-field, sine-wave gratings (0.1 cycles per degree; 2 Hz; 50% contrast) were randomly interleaved at 16 directions (22.5 deg increments) for 2 s. Stimuli alternated with a 4 s ITI of uniform mean luminance.

### Registration, segmentation, matching across sessions, and time course extraction

To adjust for x-y motion, we registered all frames from each imaging session to a stable reference image. For each experiment, we first segmented cells in the D1 session and then used this as a reference to find matching cells in the D3 and D7 sessions. Cell bodies were manually segmented by selecting all visible cells from images of the average dF/F during stimulus presentation (where F is the average of 1 s preceding each stimulus) for each unique stimulus condition.

We then found matching cells in the other two sessions by using salient fiduciary marks (e.g. bright cells and thin vasculature) to align the D3 or D7 image stack to the D1 session. Then, for each cell segmented in the D1 session we examined the corresponding region of the stack from the D3 or D7 session to determine whether the matching cell was detectable (based on location and morphological similarity). In addition, we used dsRed fluorescence to identify cells that express the gRNA, and only these cells were considered for further analysis.

Fluorescence time courses were derived by averaging all pixels in a cell mask. To exclude signal from the neuropil, we first selected a three pixel shell around each neuron (excluding a three pixel boundary around the segmented neuron and the territory of neighboring neurons), then estimated the neuropil scaling factor by maximizing the skew of the resulting subtraction and finally subtracted this component from each cell’s time course.

### Visual responses and cell inclusion

Visually-evoked responses were measured as the average dF/F during the 2 s stimulus period. Cells were included if they were identified on D1 and at least one other session and if they were visually responsive on D1 (in response to at least one stimulus condition as defined by a Bonferroni corrected paired t-test).

To measure orientation preference, responses to opposite directions were averaged, and the data was fit with a Von Mises function:

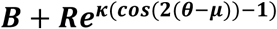

where B is the baseline firing rate, R is the modulation rate, κ is the concentration, and μ is the preferred orientation. κ is equivalent to the variance (σ2) of the fit, and is converted to standard deviation (σ) in degrees:

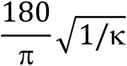

To measure the reliability of the fit, the fit was repeated 1000 times resampling trials with replacement. Reliability was measured as the 90th percentile of the difference in bootstrapped measures of preferred orientation from the original measure (i.e. a value of 20 means that 90% of bootstraps are within 20 degrees of the original fit). Cells that had a reliability of less than 22.5 degrees (i.e. the interval in our stimulus set) on D1 were considered tuned and included for measuring changes in orientation preference across sessions.

### Conditioned place prefere nce (CPP)

Behavior experiments were collected from male and female mice. Prior to acclimation day, mice were housed in the testing room at least 1 day to habituate to the room. For day 1, mice were placed into the 3-chamber box (Med Associates) and allowed to freely explore all three chambers for 30 minutes. Time spent in each box was measured through breaks in infrared beams placed in each box. For the next six days, mice were given alternating intraperitoneal (IP) injections of either saline or cocaine and placed in either the black or white box. The mice spent 30 minutes in that chamber, with infrared beambreaks used as a proxy for movement. On day 8, the mice were allowed to explore all three chambers again for 30 minutes to test for box preferences. For the initial pilot experiment, interleaved 10 minute test days were added after each cocaine injection day. For the main experiment in Figure 8, an extra acclimation day and break day were added, and the first training day was cocaine. For the pooled data in Figure 8, data came from all three versions of the paradigm as described above. CPP score was calculated with this formula:

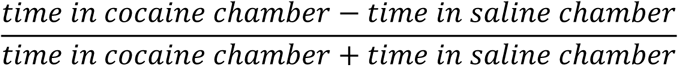

In this formula, -1 corresponds to a maximal preference for the saline side, and +1 corresponds to a maximal preference for the cocaine side.

### Prepulse inhibition

Prepulse inhibition was evaluated using the SR-LAB startle response system and associated software (San Diego Instruments, San Diego, CA). The experimental protocol comprised three trial types: null trials (64 dB white-noise background, no stimulus), pulse-alone trials (40 ms, 120 dB white-noise stimulus), and prepulse+pulse trials. In the latter, the 120 dB startle stimulus was preceded by 100 ms by a 20 ms prepulse set at 4, 8, or 12 dB above the background noise. Prior to testing, mice were placed in plexiglass cylinders and allowed to acclimate to the apparatus for five minutes. Each session consisted of 42 trials: 18 pulse-alone, 6 null (no stimulus), and 18 prepulse-pulse trials. A standardized trial order was applied to all subjects, beginning with 6 pulse-alone trials, followed by a pseudorandomized combination of prepulse+pulse trials, pulse-alone trials, and null trials, and concluding with 6 pulse-alone trials. Startle magnitude was recorded as the first maximum platform displacement (mAmp) by the animal (e.g., “startle”) within 100ms following the onset of the pulse stimulus, onset of the null trials.

PPI was quantified using the following formula: % PPI = [(Pulse Alone - Prepulse+Pulse) / Pulse Alone] × 100.

### Novel object recognition

Short-term memory (STM) and long-term memory (LTM) were assessed using the novel object recognition (NOR) task. Briefly, mice were acclimated to the empty test arena (22 x 22 x 14 cm; illuminated at 125 lux) 60 min before training. For object training the mice were reintroduced to the test chamber and allowed free exploration of an identical pair of Lego objects (8 x 2.5 x 2 cm) for 5 minutes. In the STM test (30 minutes after training), one of the familiar training objects was replaced with a novel object of similar size but different shape and features and the mouse was given 5 min free exploration. Twenty-four hours after training, the remaining familiar training object was paired with a different novel object of similar volume but new features for the LTM test.

Behaviors were video-recorded with 3-point body detection (nose-center-tail) using Noldus Ethovision 17 (Leesburg, VA). Object recognition scores were calculated by subtracting the time spent with the novel object from the time spent with the familiar object, and dividing this difference by the total amount of time spent with both objects. Total time spent with the objects, number of contacts and the activity (cm) of the animals in each test was also scored.

### Experimental Design and Statistical Analyses

Statistical tests were primarily performed in GraphPad Prism, with analyses for two-photon imaging performed using custom code written in MATLAB (Mathworks). N values refer to number of cells or mice. P < 0.05 was considered statistically significant, and significance is indicated as follows: *p < 0.05, **p < 0.01, ***p<0.001, ****p<0.0001.

## Results

### Engineered dCas3 fusion proteins permit bidirectional modulation of Arc transcription from specific gene regulatory elements

Transcription of the *Arc* gene is controlled not only by its proximal promoter, but also by a distal synaptic activity-regulated enhancer (SARE) that contains binding sites for the activity-regulated transcription factors CREB, SRF, and MEF2 (Kawashima et al., 2009). To determine how these regulatory elements contribute to the basal and stimulus-induced transcription of *Arc*, we designed guide RNAs (gRNAs) to recruit an engineered dead Cas9 (dCas9) activator (dCas9-VP64) or repressor (dCas9-KRAB) to the promoter or distal enhancer of *Arc*. As a control, we used a non-targeting gRNA directed against the sequence of the bacterial gene *LacZ*. To study stimulus-regulated transcription of *Arc,* we treated cultured embryonic cortical neurons with 50ng/ml Brain-Derived Neurotrophic Factor (BDNF), which induces robust transcription of *Arc* mRNA within minutes (El-Sayed et al., 2011). This stimulus also induces bidirectional expression of non-coding enhancer RNAs (eRNAs) from the upstream *Arc* enhancer, showing that this regulatory element is recruiting active RNA polymerase under these conditions (Schaukowitch et al., 2014).

Lentiviral infection of neurons with the dCas9-KRAB repressor together with gRNAs targeting either the enhancer or the promoter blocked the induction of *Arc* mRNA by BDNF compared with cells expressing a control gRNA (**Figure 1A**; two-way ANOVA, main effect of time: F_(2,26)_ = 388.8 p<0.0001; main effect of gRNA F_(2,26)_ = 326.6 p<0.0001; gRNA x time interaction: F_(4,26)_ = 142.1, p<0.0001; Dunnett’s post hoc, Prom 30 min, Prom 60 min, Enh 30 min, Enh 60 min all p <0.0001). Although there were main effects of time and gRNA, the pairwise comparisons show that recruitment of the dCas9-VP64 activator to the *Arc* promoter or enhancer did not further increase stimulus-dependent activation of *Arc* (**Figure 1B**; two-way ANOVA, main effect of time: F_(2,27)_ = 180.7, p<0.0001; main effect of gRNA F_(2,27)_ = 6.81, p=0.0040; gRNA x time interaction: F_(4,27)_ = 2.801, p=0.0457). This is consistent with prior studies, which show that endogenous neuronal activity-induced transcriptional mechanisms of gene activation are stronger than those of engineered dCas9 activators (Avarlaid et al., 2024; Chen et al., 2019). The effects of dCas9 recruitment were specific to *Arc* and did not reflect a general change of BDNF-induced transcription, as there was no effect of dCas9 or gRNA expression on *Fos* expression or induction in the same samples (**Figure 1C, D**; two-way ANOVA, main effect of time: F_(2,26)_ = 48.02, p<0.0001; no main effect of gRNA: F_(2,26)_ =1.406, p=0.2632; time x gRNA interaction: F_(4,26)_ = 0.5561, p=0.6964).

**Figure 1.**
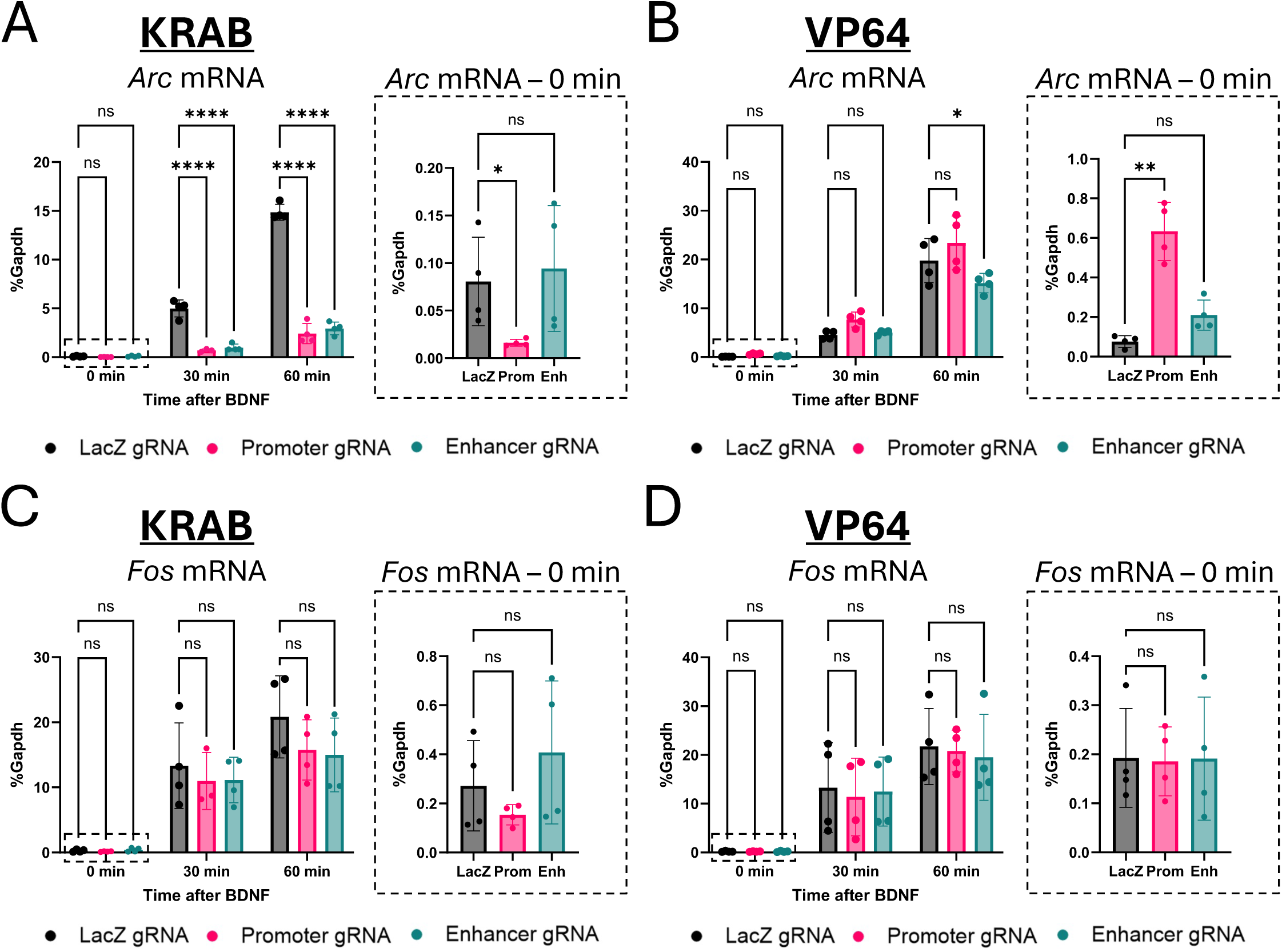
Targeted modulation of *Arc* promoter and enhancer leads to changes in mRNA expression in vitro. **(A, B)** qPCR of *Arc* mRNA from DIV7 cultured cortical neurons after targeting the *Arc* promoter or enhancer with dCas9-KRAB **(A)** or dCas9-VP64 **(B)** lentiviruses. Samples were treated with BDNF for 0, 30, or 60 minutes. Inset graphs show separate one-way ANOVAs of basal mRNA (0 min stimulation) expression. **(C, D)** Same as (A, B) for *Fos* mRNA from same samples to show specificity of manipulation to the *Arc* gene. *p<0.05, **p<0.01, ****p<0.0001. Data are represented as mean ± SD.

Notably, recruitment of dCas9-KRAB to the *Arc* promoter, but not the enhancer, reduced *Arc* expression under basal conditions (**Figure 1A** inset graph; Kruskal-Wallis H(2) = 7.385, p = 0.0145; Dunn’s post hoc test *LacZ* vs Prom p=0.0372, *LacZ* vs Enh p>0.9999). Similarly, recruitment of dCas9-VP64 to the *Arc* promoter, but not the enhancer, significantly increased *Arc* mRNA under basal conditions prior to BDNF addition (**Figure 1B** inset graph; Kruskal-Wallis H(2) = 9.846, p = 0.0002; Dunn’s post hoc test *LacZ* vs Prom p=0.0034, *LacZ* vs Enh p=0.2333). Importantly, there was no effect of dCas9-KRAB or dCas9-VP64 manipulation on baseline *Fos* expression (**Figure 1C, D** inset graphs; KRAB: Kruskal-Wallis H(2) = 2.000, p=0.3967; VP64: Kruskal-Wallis H(2) = 0.0385, p=0.9945).

To confirm that the enhancer-targeted gRNAs could modulate local transcription at the enhancer, we measured expression of the *Arc* eRNAs in the same samples used for **Figure 1**. dCas9-KRAB blocked BDNF-induced *Arc* positive strand (**Figure 2A**; two-way ANOVA, main effect of time: F_(2,25)_ = 29.39, p<0.0001; main effect of gRNA: F_(2,25)_ = 26.49, p<0.0001; gRNA x time interaction: F_(4,25)_ = 5.622, p=0.0023; Dunnett’s post hoc, Enh 30 min, Enh 60 min p <0.0001) and negative strand (**Figure 2C**; two-way ANOVA, main effect of time: F_(2,26)_ = 39.74, p<0.0001; main effect of gRNA: F_(2,26)_ = 28.50, p<0.0001; gRNA x time interaction: F_(4,26)_ = 7.287, p=0.0004; Dunnett’s post hoc, Enh 30 min, Enh 60 min p <0.0001) eRNA expression only when it was recruited to the enhancer, not the promoter. Similar to its effects on *Arc* mRNA, expression of dCas9-VP64 had no effect on stimulus-induced *Arc* positive strand eRNA expression (**Figure 2B**; two-way ANOVA, main effect of time: F_(2,27)_ = 35.08, p<0.0001; main effect of gRNA: F_(2,27)_ = 0.005, p=0.9951; gRNA x time interaction: F_(4,27)_ = 0.3541, p=0.8389) but did have mild effects on negative strand eRNA expression (**Figure 2D**; two-way ANOVA, main effect of time: F_(2,27)_ = 72.91, p<0.0001; main effect of gRNA: F_(2,27)_ = 9.472, p=0.0008; gRNA x time interaction: F_(4,27)_ = 0.2992, p=0.8759; Dunnett’s post hoc, Enh 30 min, p=0.0325).

**Figure 2.**
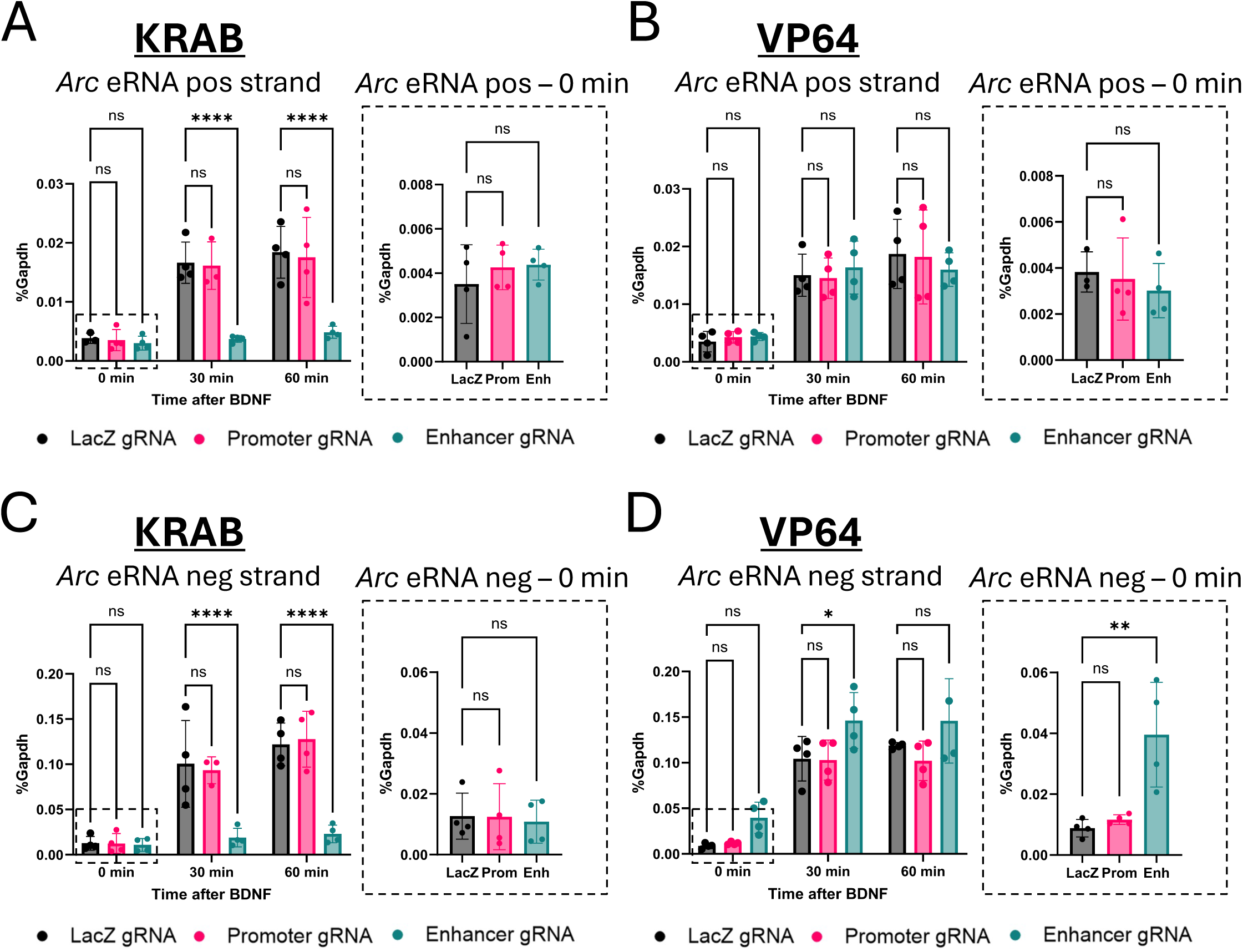
Targeted modulation of *Arc* promoter and enhancer leads to changes in enhancer RNA (eRNA) expression in vitro. **(A, B)** qPCR of *Arc* pos-strand eRNA from DIV7 cultured cortical neurons after targeting the *Arc* promoter or enhancer with dCas9-KRAB **(A)** or dCas9-VP64 **(B)** lentiviruses. Samples were treated with BDNF for 0, 30, or 60 minutes. Inset graphs show separate one-way ANOVAs of basal eRNA (0 min stimulation). **(C, D)**. Same as (A, B) for *Arc* neg-strand eRNA. *p<0.05, **p<0.01, ****p<0.0001. Data are represented as mean ± SD.

Under basal conditions there was minimal *Arc* eRNA expression and no effect of dCas9-KRAB at either the promoter or the enhancer on positive strand (**Figure 2A** inset graph; Kruskal-Wallis H(2) = 0.7308, p = 0.7463) or negative strand (**Figure 2C** inset graph; Kruskal-Wallis H(2) = 0.2692, p = 0.9131) eRNA. Although dCas9-VP64 recruitment to the enhancer did not cause a significant basal increase in positive strand *Arc* eRNA (**Figure 2B** inset graph; Kruskal-Wallis H(2) = 2.053, p = 0.3879), it did increase the expression of the negative strand (**Figure 2D** inset graph; Kruskal-Wallis H(2) = 8.346, p = 0.0024; Dunn’s post hoc *LacZ* vs Enh 30 p=0.0089).

Taken together, these data show that by using dCas9 fused to activators or repressors, we can modulate *Arc* transcription via recruitment either to the proximal promoter or the distal enhancer. Further, these data suggest that whereas activation of the promoter is required for *Arc* transcription under both basal and activity-induced conditions, the enhancer is selectively engaged to regulate *Arc* transcription only in response to cellular stimulation.

### dCas3-regulated transcription of Arc is sufficient to change levels of Arc protein

In addition to being controlled at the transcriptional level, Arc protein levels are also determined by stimulus-dependent translational regulation in neurons (Das et al., 2023; Giorgi et al., 2007). To determine whether transcriptional modulation of *Arc* using engineered dCas9 activators and repressors is sufficient to change Arc proteins levels, we harvested BDNF-treated dCas9-expressing cortical neurons to visualize Arc protein expression by western. Similar to our data for *Arc* mRNA, we found that recruitment of dCas9-KRAB to either the promoter or the enhancer of *Arc* was sufficient to block BDNF-induced Arc protein accumulation in neurons (**Extended data Figure 1A**). Arc protein expression was too low in these embryonic neuron cultures under basal conditions to determine by western whether there were any significant effects of promoter or enhancer repression prior to BDNF treatment. Recruitment of dCas9-VP64 to the *Arc* promoter increased expression of Arc both under basal conditions and following BDNF stimulation, whereas there were no increases following artificial activation of the *Arc* enhancer (**Extended data Figure 1B**). Immunocytochemistry of neuronal coverslips qualitatively show similar patterns (**Extended data Figure 1C, D**). Taken together, these data show that we can use dCas9 fusion proteins targeted to either the promoter or the enhancer of the *Arc* gene to determine the functional consequences of stimulus-regulated Arc expression in neuronal plasticity.

### Transgenic dCas3-KRAB mice permit regulation of Arc protein induction in response to light in vivo

To determine whether dCas9-KRAB regulation of Arc transcription is sufficient to block induction of Arc protein in response to physiologically-relevant stimuli in vivo, we injected visual area V1 of adult dCas9-KRAB transgenic mice (Gemberling et al., 2021) with AAVs that expressed Cre recombinase and a fluorophore together with gRNAs targeting the *Arc* promoter, the *Arc* enhancer, or *LacZ* as a control. Mice were dark housed for 4 days and brains were either harvested immediately after dark housing or after 4 hours of light exposure, which robustly induces Arc expression in V1 (Wang et al., 2006). Because *Arc* expression is regulated by sensory experience, which can cause Arc to vary significantly between individual mice, we conducted within-mouse controls, delivering *LacZ* gRNAs into the left hemisphere, and either *Arc* promoter or *Arc* enhancer gRNAs into the right hemisphere (**Figure 3A, B**).

**Figure 3.**
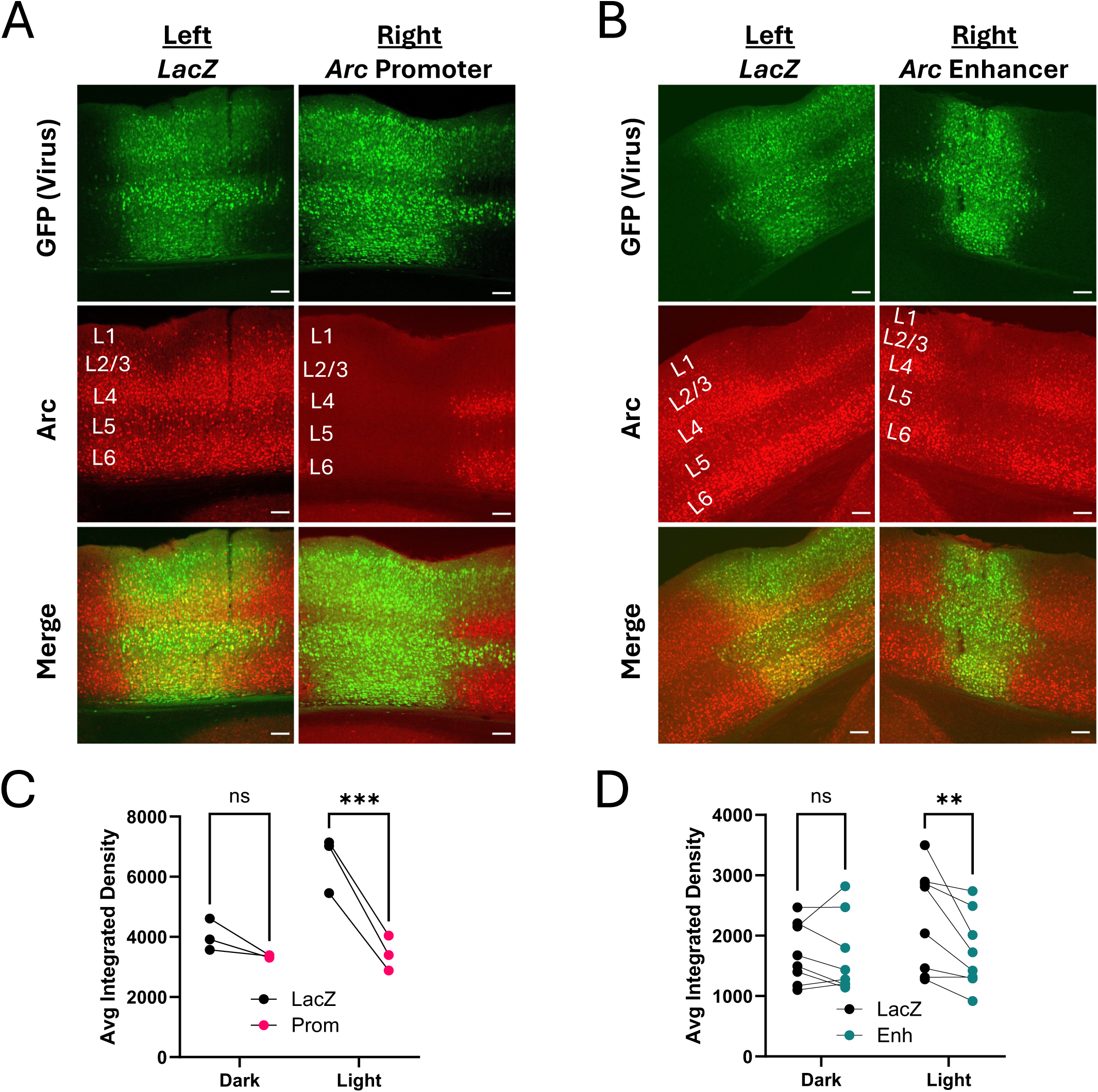
Arc knockdown in visual cortex after targeted repression of either the *Arc* promoter or enhancer. **(A)** 10x images of mouse V1 showing strong Arc knockdown through dCas9-KRAB targeted to the *Arc* promoter. **(B)** Arc knockdown still occurred through targeting the *Arc* enhancer but to a lesser extent. Scale bars are 100μm. **(C)** Within-mouse comparisons of average cell integrated densities from left (*LacZ* gRNA) and right (*Arc* promoter) hemispheres showing Arc knockdown from promoter repression. Cells from all layers were quantified. Dark mice were perfused immediately following four days of dark housing, and light mice were perfused after four hours of light exposure. **(D)** Same as (C) except right hemisphere targeted the *Arc* enhancer and layer 6 was not quantified due to imaging constraints. **p<0.01, ***p<0.001.

When compared with the *LacZ*-expressing hemisphere, Arc expression was significantly decreased when dCas9-KRAB was targeted to either the *Arc* promoter (**Figure 3C**, n = 3 mice per condition; two-way rmANOVA, main effect of gRNA: F_(1,4)_ = 79.12, p=0.0009; main effect of light: F_(1,4)_ = 8.132, p=0.0463; gRNA x light interaction: F_(1,4)_ = 32.59, p=0.0047; Light: *LacZ* vs Enh p=0.0010) or the *Arc* enhancer (**Figure 3D**, n = 6 mice per condition; two-way rmANOVA, main effect of gRNA: F_(1,14)_ = 6.812, p=0.0206; main effect of light: F_(1,14)_ = 0.9836, p=0.3382; gRNA x light interaction: F_(1,4)_ = 4.991, p=0.0423; Light: *LacZ* vs Enh p=0.0041). These data show that inhibition of *Arc* transcription is sufficient to reduce the light-induced levels of Arc protein in visual cortical neurons.

Previously, we also reported our ability to use another transgenic mouse line carrying a Cre-inducible dCas9-p300 transgene to modulate expression of *Fos* mRNA and protein in hippocampal neurons *in vivo* (Gemberling et al., 2021). To determine whether we could use this mouse line to elevate Arc expression in visual cortex, we injected control *LacZ* gRNAs or gRNAs targeting the *Arc* promoter into visual cortex area V1 as we had done for the dCas9-KRAB mice. However, when we examined the region of V1 that had viral expression as indicated by the viral fluorophore, we saw a striking loss, not a gain, of Arc protein expression, and this was true even in the sections expressing only the *LacZ* control gRNA (**Figure 4A**). Further, in these mice, Fos expression was also reduced in virally expressing cells, suggesting there was a general decrease in activity-dependent gene expression. Although expression of dCas9-p300 can drive increases in histone acetylation at lysine 27 (H3K27ac)(Gemberling et al., 2021) we were surprised to find reduced H3K27ac in virally infected cells in the dCas9-p300 mice (**Figure 4A**). These cells also showed reduced expression of NeuN, suggesting a global impact on neuronal transcription. These changes were not seen when comparing the virally-infected region of dCas9-KRAB mice expressing the *LacZ* gRNA to their uninfected surround, indicating this was specific to the dCas9-p300 transgene, and not a general effect of expressing dCas9 expression (**Figure 4B**). Notably, a recent study reported gRNA-independent changes in striatal behaviors when the Cre-inducible dCas9-p300 strain was crossed to D1-Cre but not D2-Cre transgenic mice (Campbell et al., 2025). These data suggest that long-term in vivo expression of the active enzymatic domain of p300 in specific cell types (e.g. visual cortical neurons, D1 spiny projection neurons) of the dCas9-p300 mice may induce cellular compensations that limit the utility of this transgene to control gene expression in vivo. Given these concerns, the remainder of the work focuses on CRISPRi through dCas9-KRAB.

**Figure 4.**
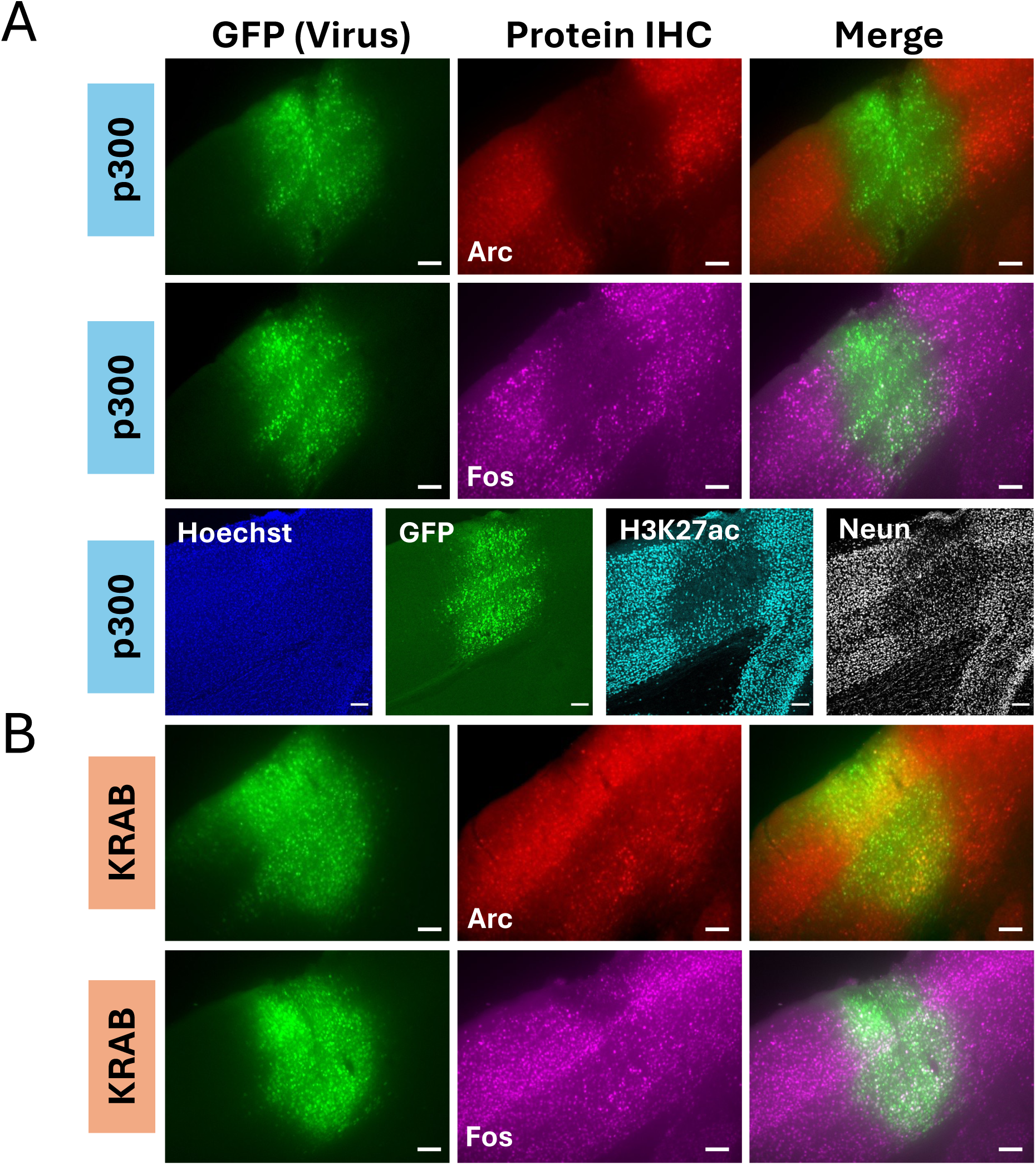
Nonspecific effects of transgenic dCasG-p300 on gene expression in the visual cortex. **(A)** 10x images of visual cortex from a dCas9-p300 mouse that was injected with a control *LacZ* gRNA virus. Slices were stained for Arc, Fos, H3K27ac, or NeuN to look for nonspecific effects of transgenic expression of dCas9-p300. **(B)** 10x image of a dCas9-KRAB mouse with *LacZ* gRNA virus as a control to show intact Arc expression in infected cells. Scale bars are 100μm.

### CRISPR inhibition of Arc in V1 has a limited effect on orientation selectivity at baseline and neither enhances nor impairs drift of orientation tuning over days

Prior studies have shown that Arc is required during development for the experience-dependent refinement of visual cortex connectivity that establishes the number of binocular neurons (Jenks & Shepherd, 2020). Knocking down Arc in adult mouse visual cortex also increases the number of neurons that show binocular responses (Jenks & Shepherd, 2020). However, unlike in other brain regions, like the dentate gyrus of the hippocampus (**Extended data Figure 2**), levels of Arc expression in adult cortex are high at baseline (Jenks et al., 2017), and whether experience-dependent increases in levels of Arc expression contribute to adult plasticity is not well understood. Interestingly, a recent study showed that Arc is among a set of activity-regulated genes that are induced daily in visual cortex of mice in concert with the light/dark cycle (Kushinsky et al., 2024). Knockdown of one of the other daily light-regulated genes, *Npas4*, was shown to disrupt the stability of orientation tuning. Thus, we decided to use calcium imaging of visual cortical neuron responses over days to determine the requirement for the experience-dependent regulation of Arc in plasticity of adult visual cortex.

To test whether CRISPRi of Arc expression impacts response properties of visual cortical neurons, we targeted adeno-associated viral (AAV) injections to V1 of dCas9-KRAB transgenic mice to express the genetically encoded calcium indicator GCaMP7f along with a second AAV that expressed Cre recombinase, nuclear-targeted dsRed, and a gRNA (n = 14 mice, 8 males, 6 females: 7 *LacZ*, 4 *Arc* Promoter, 3 *Arc* Enhancer). To measure orientation selectivity and the stability of this feature in single neurons over time, we used two-photon imaging to measure visually-evoked responses from populations of neurons in layer 2/3 in response to sinusoidal drifting gratings at 8 orientations (0.1 cycles/degree, 2 Hz, 50% contrast) and used vascular landmarks to return to the same population of neurons for three imaging sessions (Day 1 (D1), D3 and D7; **Figure 5A, B**).

**Figure 5.**
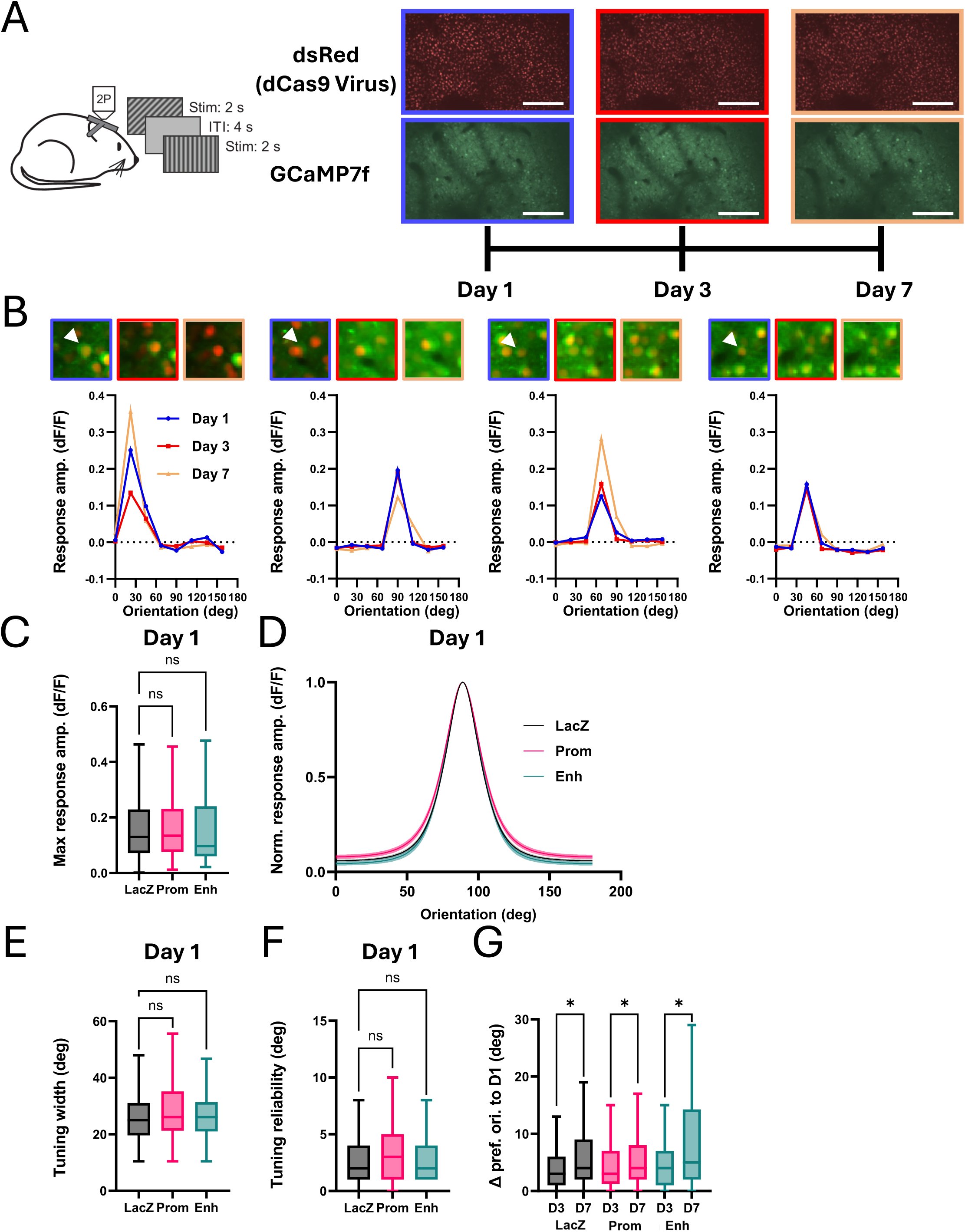
Arc knockdown in visual cortex does not strongly affect orientation tuning or stability of representations. **(A)** Schematic of experimental paradigm measuring orientation tuning across multiple sessions. The same neurons were imaged on Days 1 (blue), 3 (red), and 7 (orange). Scale bars are 100μm. **(B)** Orientation tuning curves from four example cells imaged across the three sessions in the experiment in (A). White arrow in postage stamp images points to cell being measured. **(C)** Maximum normalized change in fluorescence (dF/F) across orientations for each cell on the first imaging session for mice expressing *LacZ* (black; n = 333 cells, 7 mice), *Arc* Promoter (pink; n = 173 cells, 3 mice) or *Arc* Enhancer (teal; n = 94 cells, 3 mice) gRNA. **(D)** Average orientation tuning curve from Von Mises fit. Error is SEM across cells. **(E, F)** Same as (C) for the sharpness of tuning and reliability of fit. **(G)** Change in preferred orientation for cells compared on Days 1 and 3 [LacZ (black; n = 272 cells, 7 mice), *Arc* Promoter (pink; n = 128 cells, 4 mice) or *Arc* Enhancer (teal; n = 82 cells, 3 mice)] and Days 1 and 7 [LacZ (black; n = 233 cells, 7 mice), *Arc* Promoter (pink; n = 110 cells, 4 mice) or *Arc* Enhancer (teal; n = 68 cells, 3 mice)]. Two-way ANOVA reveals main effect of interval between imaging sessions (p = 0.003) but no main effect of Arc manipulation (p = 0.12). *p<0.05. Box plots reflect the median and 25th to 75th percentiles; outliers are not presented. Whiskers are 1.5x IQR (Tukey).

We first assessed neuronal responses in a single imaging session of the population of neurons that co-express dsRed (and therefore dCas9-KRAB and the gRNA) and GCaMP7f that were significantly responsive to at least one visual stimulus condition on the first day of imaging (D1: *LacZ* – n = 333 cells; *Arc* Promoter – n = 173 cells; *Arc* Enhancer – n = 94 cells). We found no significant difference in the amplitude of the response to the preferred orientation across conditions (**Figure 5C**; one-way ANOVA, F_(2, 597)_ = 0.087, p=0.9167). There is a small but significant broadening of orientation tuning upon Arc manipulation; however, this effect was not strong enough to show significance in individual post hoc comparisons (**Figure 5D, E**; one-way ANOVA, F_(2, 597)_ = 2.421, p=0.0897; Tukey’s post hoc comparisons: *LacZ* vs *Arc* Promoter, p=0.0545; *LacZ* vs *Arc* Enhancer, p=0.7881). We also found no difference in the reliability of orientation tuning (**Figure 5F**; one-way ANOVA, F_(2, 597)_ = 1.082, p=0.3397), suggesting that there is no change in the variability of responses across trials.

Notably, the reliability across all cells was extremely high, with 89% of cells having less than 10 degrees of variation of preferred orientation within session. Thus, we find that CRISPRi of Arc expression has measurable but limited effects on the orientation tuning of neurons in V1 of adult mice.

We next investigated the stability of orientation tuning across sessions. For these analyses, we considered the subset of neurons that were both well-tuned for orientation (reliability < 22.5 deg) and could be identified across multiple imaging sessions (D1 vs D3: *LacZ* – n = 272 cells; *Arc* Promoter – n = 128 cells; *Arc* Enhancer – n = 82 cells; D1 vs D7: *LacZ* – n = 233 cells; *Arc* Promoter – n = 110 cells; *Arc* Enhancer – n = 68 cells), and measured the change in preferred orientation across sessions. Similar to the within-session reliability, we see a high degree of reliability across sessions with 83% of cells having less than 10 degrees change in preferred orientation between D1 and D3, and 75% having less than 10 degrees change between D1 and D7. This larger change in orientation preference between sessions D1 and D7 than between D1 and D3 is significant (**Figure 5G**, two-way rmANOVA: main effect of interval, F_(1, 889)_ = 9.05, p=0.0027; *LacZ* D3 vs D7: p=0.0314, Prom D3 vs D7 p=0.0314, Enh D3 vs D7 p=0.0314) and consistent with previous work into the dynamics of representational drift (Bauer et al., 2024). However, we find no significant effect of manipulation of Arc on this plasticity of orientation preference (main effect of gRNA, F_(2, 889)_ = 2.14; p = 0.1186). Thus, while we do see evidence for representational drift of orientation selectivity in area V1, Arc does not seem to be important for either the plasticity or the stability of this form of activity-dependent cortical plasticity.

### CRISPR knockdown of Arc in the nucleus accumbens does not block cocaine-induced place preference

Although our calcium imaging data support the presence of some ongoing plasticity in the primary visual cortex of adult mice, it is the subcortical structures that are thought to be most sensitive to environmental adaptations in the context of learning and memory as well as reward-triggered behaviors. These regions have more sparse, ensemble-like expression of Arc at baseline compared with cortex (**Extended data Figure 2**), and thus we suspected they might be more likely to reveal stimulus-induced Arc functions. Cocaine is known to induce synapse plasticity of spiny projection neurons in the nucleus accumbens (NAc)(Dong & Nestler, 2014). Experimental manipulations of synapse strength in NAc can modulate behaviors induced by cocaine (Lee et al., 2013), and at least two studies using conditional knockdown strategies previously implicated Arc in some aspects of cocaine-triggered behaviors (Barry et al., 2023; Wood et al., 2024). Thus, we next asked whether dCas9-KRAB mediated knockdown of Arc in neurons of the NAc would change circuit or behavioral responses to cocaine.

As has been previously reported (Salery et al., 2017), acute cocaine exposure drives induction of Arc expression within 2 hours in the core and shell of the NAc. We observe both bright signal in the nuclei of NAc neurons, and a hazy signal throughout the NAc that likely represents Arc protein in the dendrites of these neurons (**Figure 6**). To determine whether dCas9-KRAB recruitment to the *Arc* promoter or enhancer is sufficient to block Arc protein induction in NAc by cocaine exposure in vivo, we again conducted within-animal control studies. We stereotaxically injected *LacZ* gRNA viruses into the left NAc and either *Arc* promoter or enhancer targeting gRNAs into the right NAc of the same dCas9-KRAB transgenic mice. Three weeks later, mice were injected with 20mg/kg i.p. cocaine to induce Arc expression as measured by immunostaining. Targeting dCas9-KRAB to either the *Arc* promoter or the *Arc* enhancer significantly reduced the number of Arc+ neurons in NAc (**Figure 6A, B**; n = 4-5 mice; paired two-tailed t-test, *LacZ* vs Prom p=0.0045, *LacZ* vs Enh p=0.0250). By contrast, Fos was still induced in the regions of the NAc that expressed the virus and lacked Arc (**Figure 6C, D**; n = 4-5 mice; paired two-tailed t-test, *LacZ* vs Prom p=0.7256, *LacZ* vs Enh p=0.0687. Thus, these data show that CRISPRi is sufficient to selectively block Arc induction by cocaine in the NAc.

**Figure 6.**
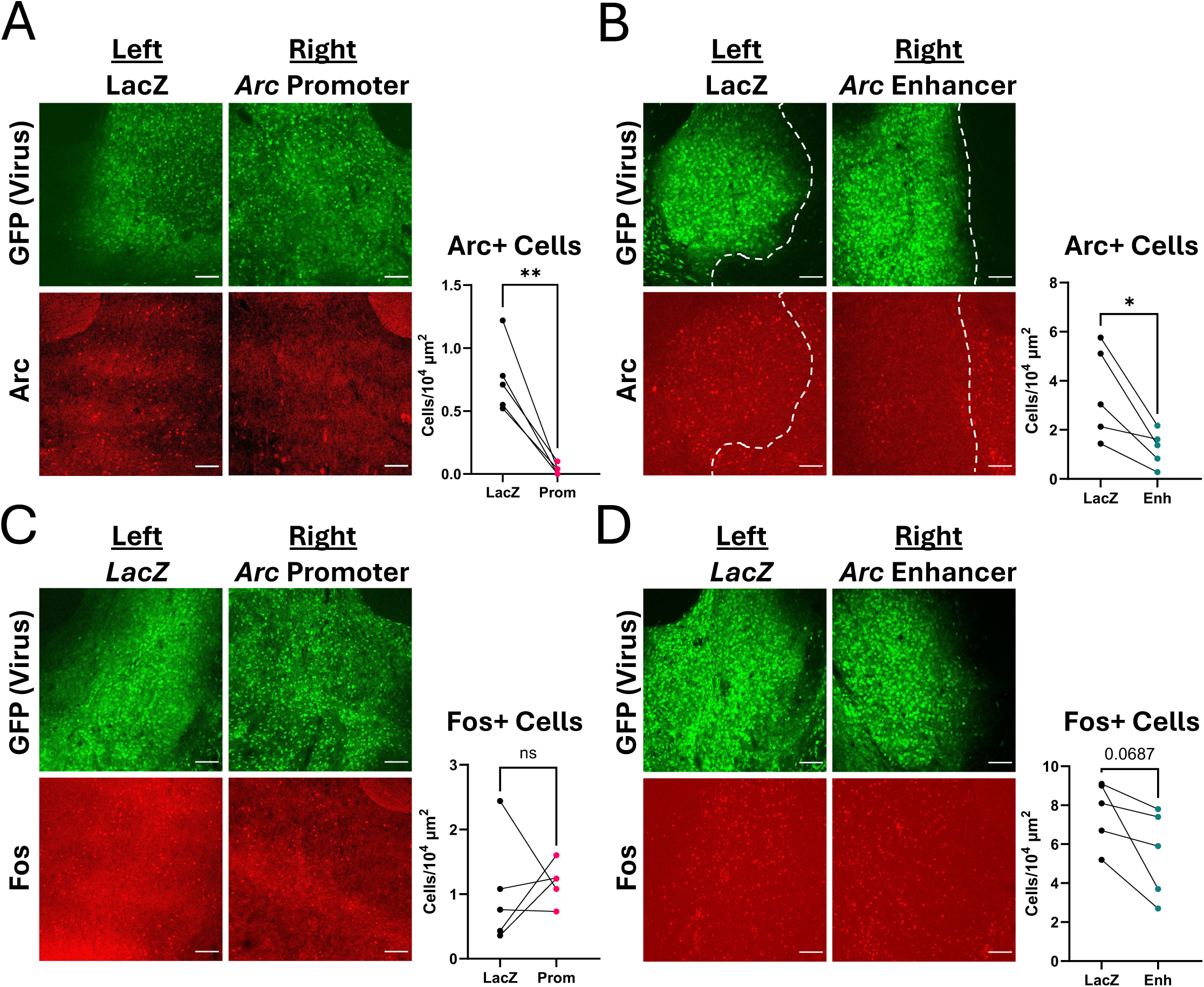
Arc knockdown in the nucleus accumbens (NAc) through promoter or enhancer targeting. **(A, B)** dCas9-KRAB mice were injected with a control *LacZ* gRNA virus (GFP) in the left NAc and either *Arc* promoter **(A)** or *Arc* enhancer **(B)** viruses in the right NAc. Mice were injected with 20mg/kg cocaine, perfused one hour later, and brains were harvested for Arc staining (red). Number of Arc+ cells were quantified. **(C, D)** Same (A, B) for Fos expression (red). Quantification was performed either by hand by blinded observer (A, C) or by Ilastik analysis software (B, D). Scale bars are 100μm. *p<0.05, **p<0.01.

To test the behavioral consequences of inhibiting the cocaine-induced expression of Arc in the NAc, we stereotaxically injected dCas9-KRAB mice bilaterally in the NAc with viruses co-expressing Cre, dsRed, and either *LacZ*, *Arc* promoter, or *Arc* enhancer gRNAs. Prior studies using a strain of *Arc* knockin mice in which GFP replaces the *Arc* gene body, (Penrod et al., 2020; Salery et al., 2017) or *Arc* shRNAs injected in NAc (Wood et al., 2024) have reported opposite or no effects (Penrod et al., 2020) of Arc loss on cocaine-induced conditioned place preference (CPP). However, the *Arc* GFP knockin mice are thought to have developmental circuit defects in the dopamine system as indicated by reduced paired-pulse inhibition (PPI) (Managò et al., 2016), which may confound interpretation of CPP studies. We importantly confirmed that startle responses and PPI are both largely unaffected by CRISPR inhibition of Arc in the NAc (**Extended data Figure 3**), allowing us to assess the local consequences of NAc Arc inhibition on cocaine-induced behavioral plasticity.

To ensure we could detect either enhanced or reduced CPP with Arc manipulation, we performed a pilot study with an intermediate dose of 10mg/kg i.p. cocaine versus saline (**Figure 7A**, n = 19 mice; 8 males, 11 females; 4-5 mice per group). Interleaved 10 minute test days were included after each pairing to assess the trajectory of CPP development, as shown previously (Hazlett et al., 2024). We recorded total locomotor activity on each day of the study to assess the pharmacological response to the drug treatment. There were no significant differences in the locomotor activity between *LacZ* control or Arc knockdown mice, either in the saline- or cocaine-treated mice (**Figure 7B**; two-way rmANOVA, main effect of gRNA: F_(3,15)_ = 1.26, p=0.324). Additionally, the cocaine-treated mice showed enhanced locomotor activity relative to saline-treated only on the third pairing and not on the first or second pairing. This suggested that this CPP paradigm is relatively weak, as we expected.

**Figure 7.**
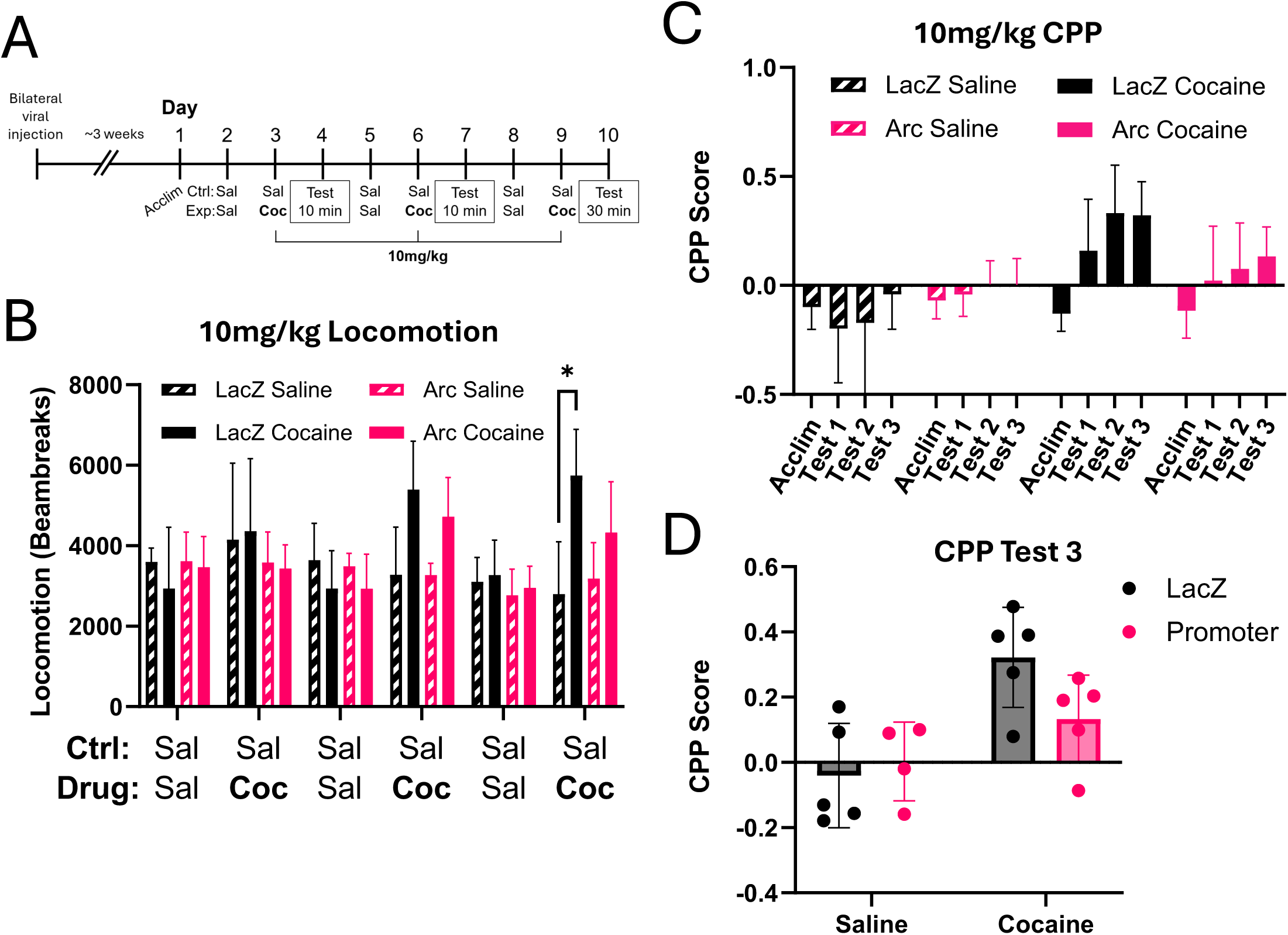
CRISPRi of the *Arc* promoter showed no difference in conditioned place preference (CPP) experiment with 10mg/kg cocaine. **(A)** Surgery and CPP timeline. dCas9-KRAB mice were bilaterally injected with *LacZ* or *Arc* promoter viruses into the NAc. After allowing for viral expression, mice were tested in a CPP paradigm with three injections of 10mg/kg cocaine and compared to control mice only receiving saline. **(B)** Locomotion of mice across days based on beambreaks measured in behavioral boxes. **(C)** CPP score across acclimation and three testing days. CPP score calculated as (time in cocaine box – time in saline box) ÷ (time in cocaine box + time in saline box). Mice were assigned boxes biased to their original preference in acclimation, which gives all mice an initial negative CPP score for acclimation. **(D)** Scores from test 3 (30 minutes) show a main effect of treatment but not of gRNA in two-way ANOVA. *p<0.05.

**Figure 8.**
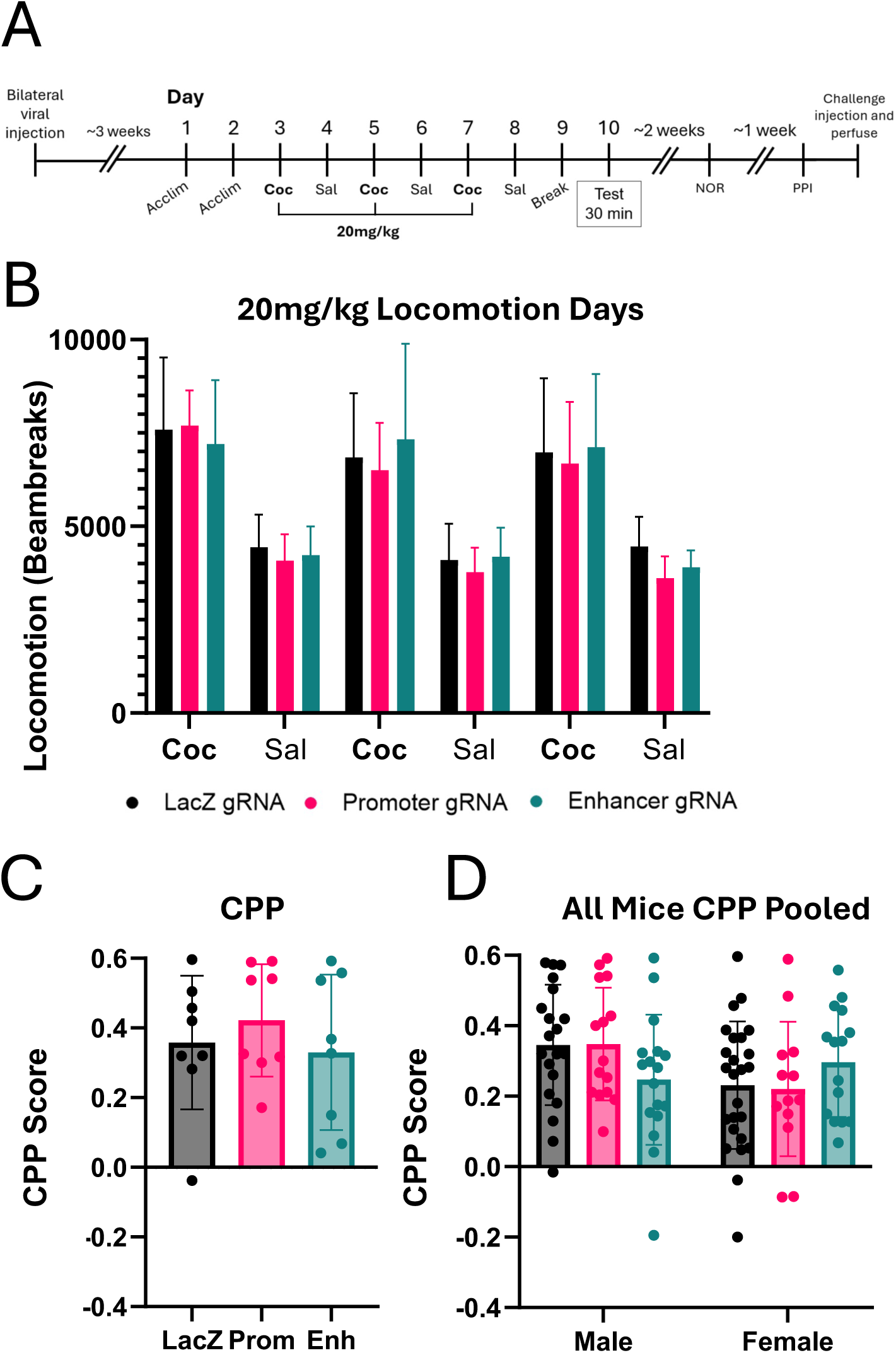
CRISPRi of *Arc* promotor or enhancer showed no difference in conditioned place preference (CPP) with 20mg/kg cocaine. **(A)** Surgery and CPP timeline. dCas9-KRAB mice were bilaterally injected with *LacZ*, *Arc* promoter, or *Arc* enhancer viruses into the NAc. After allowing for viral expression, mice were tested in a CPP paradigm with three injections of 20mg/kg cocaine. These mice were used in follow-up novel object recognition (NOR), prepulse inhibition (PPI), and comparative immunostaining in later figures. **(B)** Locomotion of mice across CPP days. **(C)** CPP scores for each condition. CPP score calculated as (time in cocaine box – time in saline box) ÷ (time in cocaine box + time in saline box). **(D)** Pooled CPP data from all injection experiments. All mice received 3 injections of either 10mg/kg or 20mg/kg cocaine.

**Figure 9.**
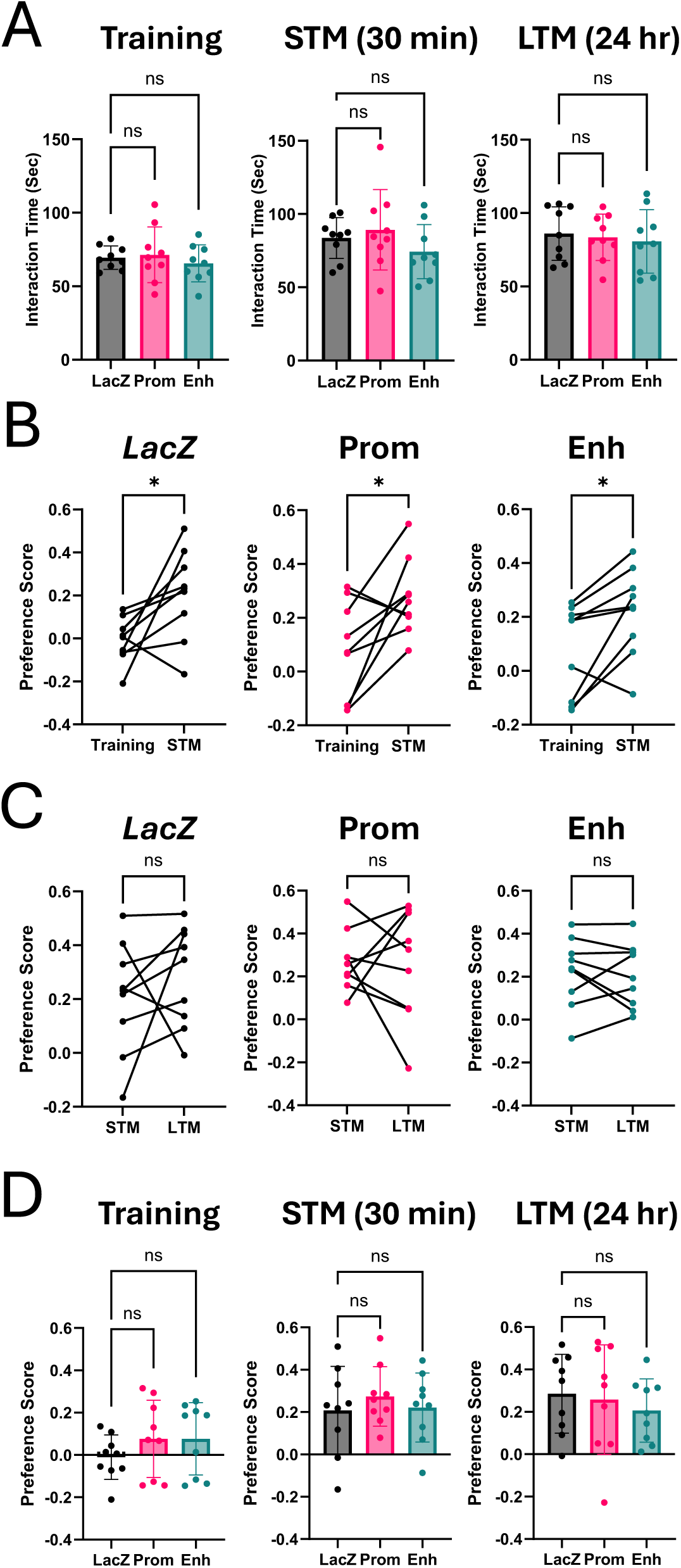
Arc knockdown in nucleus accumbens does not lead to changes in novel object recognition (NOR). dCas9-KRAB mice used were from CPP behaviors in Figure 8. **(A)** Comparisons of amount of total time spent interacting with objects for each condition in each stage of NOR. **(B)** Object preferences in training versus short-term memory (STM). Preference score calculated as (time with novel object – time with familiar object) ÷ (time with novel object + time with familiar object). **(C)** Object preferences in STM versus long-term memory (LTM). **(D)** Comparisons of preference score between conditions for all three tests. *p<0.05.

Saline-treated mice did not show a strong preference for the target chamber over the time course of the test, while both *LacZ* and *Arc* mice exposed to cocaine did increase preference over time for the cocaine-paired chamber (**Figure 7C**). However, although there appeared to be a trend of lower CPP score in the *Arc* knockdown mice on the day of the final test, there was no main effect of gRNA (**Figure 7D**; two-way ANOVA, main effect of gRNA: F_(1,15)_ = 1.200, p=0.2905; main effect of drug F_(1,15)_ = 13.69, p=0.0021; gRNA x drug interaction: F_(1,15)_ = 3.049, p=0.1012). These data suggested that the *Arc* knockdown mice might be less sensitive to the plasticity-inducing effects of cocaine, but this paradigm may be insufficient to show this effect.

We thus repeated the CPP experiment on a new cohort of mice with three pairings 20mg/kg i.p. cocaine comparing control mice with those expressing gRNAs targeting the *Arc* promoter or enhancer (**Figure 8A**, n=24; 11 males, 13 females; 8 mice per condition). We eliminated the interleaved tests to avoid any potential dampening of effect from extinction of the preference and focused solely on cocaine-conditioned mice. At this dose of cocaine, we saw robust locomotor responses on the cocaine treatment days that were not different between the three treatment groups (**Figure 8B**; two-way rmANOVA, main effect of gRNA: F_(2,21)_ = 0.2763, p=0.7613; main effect of day F_(2.360, 49.57)_ = 59.86, p<0.0001; gRNA x drug interaction: F_(4.721, 49.57)_ = 0.5080, p=0.7590). There were also no significant differences in CPP score between the three treatment groups (**Figure 8C**; one-way ANOVA, F_(2,21)_ = 0.4719, p=0.6303). In total across all of our studies using different versions of CPP, we analyzed over 100 mice and found no difference by treatment even when broken out by sex (**Figure 8D**, n=105; 52 males, 53 females; 13-25 mice per group; two-way ANOVA, main effect of gRNA: F_(2,99)_ = 0.0866, p=0.9171; main effect of sex: F_(1,99)_ = 3.337, p=0.0708; gRNA x sex interaction: F_(2,99)_ = 2.600, p=0.0793).

In addition to mediating responses to cocaine, the NAc also plays a role in gating behavioral responses to novelty (Rothwell et al., 2011). At least one previous study has shown that shRNA-mediated knockdown or Arc in the NAc impaired behavioral responses to social novelty as well as novel object interaction (Penrod et al., 2019). We noted that prior studies of Arc in the hippocampus have indicated that Arc is required for memory but not learning (Gao et al., 2018). Thus, we employed a three-step version of the novel object recognition task that starts with habituation to two identical objects and then has both a short term (30 min) as well as a long-term (24 hr) test of memory for a novel object.

All three treatment groups spent an equal amount of time exploring the two objects both during the training period and the testing periods (**Figure 9A**, one-way ANOVAs: Training F_(2, 24)_ = 0.3995, p=0.6751; STM F_(2, 24)_ = 1.189, p=0.3217; LTM F_(2, 24)_ = 0.1879, p=0.8299). When one of the objects was switched, all three groups showed a significant preference for the novel object when returned to the arena 30 min later (**Figure 9B**, paired t-tests: *LacZ* p=0.0293, Prom p=0.0260, Enh p=0.0172). Finally, when mice were returned to the arena 24 hr later, all three groups still preferred the novel object, suggesting they had no memory impairment in this specific task (**Figure 9C**, paired t-tests: *LacZ* p=0.4151, Prom p=0.8762, Enh p=0.7121). There were no differences in preference scores for any test between gRNA conditions (**Figure 9D**, one-way ANOVAs: Training F_(2, 24)_ = 0.9178, p=0.4130; STM F_(2, 24)_ = 0.3584, p=0.7024; LTM F_(2, 24)_ = 0.3591, p=0.7020). Taken together, these studies show that CRISPR-mediated inhibition of Arc induction in the adult NAc does not disrupt either novel object memory or cocaine-induced place preference.

### Inhibition of stimulus-inducible Arc expression by CRISPR inhibition of the Arc enhancer lowers the magnitude of Fos induction in NAc ensembles

In brain regions where Arc has been suggested to have a role in learning and memory, Arc is expressed in small, distributed populations of neurons known as ensembles (Josselyn et al., 2015). These neurons are thought to co-fire during learning of a task, and reactivation of the ensemble can in some cases reactivate expression of a memory (Josselyn & Tonegawa, 2020; Ramirez-Amaya et al., 2013). Immediate-early genes (IEGs) including Arc but most often Fos are used to label these ensembles (Salery et al., 2025), and in the NAc there is significant evidence that both Fos and Arc are co-expressed in the same neurons upon cocaine exposure (Thibeault et al., 2025).

To determine whether disruption of Arc induction impacts the activation of the Fos ensemble, we injected the mice previously used for the CPP and novel object studies with a challenge dose of 20mg/kg cocaine, placed them in an empty novel cage, and perfused them one hour later. We then stained for Arc or Fos on neighboring sections through the NAc and determined the number of Arc or Fos positive cells as indicated by nuclear expression above threshold using the machine learning program Ilastik. Additionally, we quantified the expression level of Arc or Fos using integrated density measurements within those positive cells.

Similar to our observations in cocaine-naïve mice (**Figure 6**), we observed that targeting the *Arc* promoter or the enhancer led to a decrease in the number of Arc+ cells induced by cocaine when compared to *LacZ* gRNA mice (**Figure 10A, C**; one-way ANOVA, F_(2, 21)_ = 8.659, p=0.0018, *LacZ* vs Prom: p=0.0032; *LacZ* vs Enh: p=0.0025). By contrast, there were no differences in the number of Fos+ cells between gRNA conditions (**Figure 10B, E**; one-way ANOVA, F_(2, 21)_ = 0.8440, p=0.4441). This shows that the CRISPR inhibition of Arc expression is specific and does not have a global effect on activity-regulated genes.

**Figure 10.**
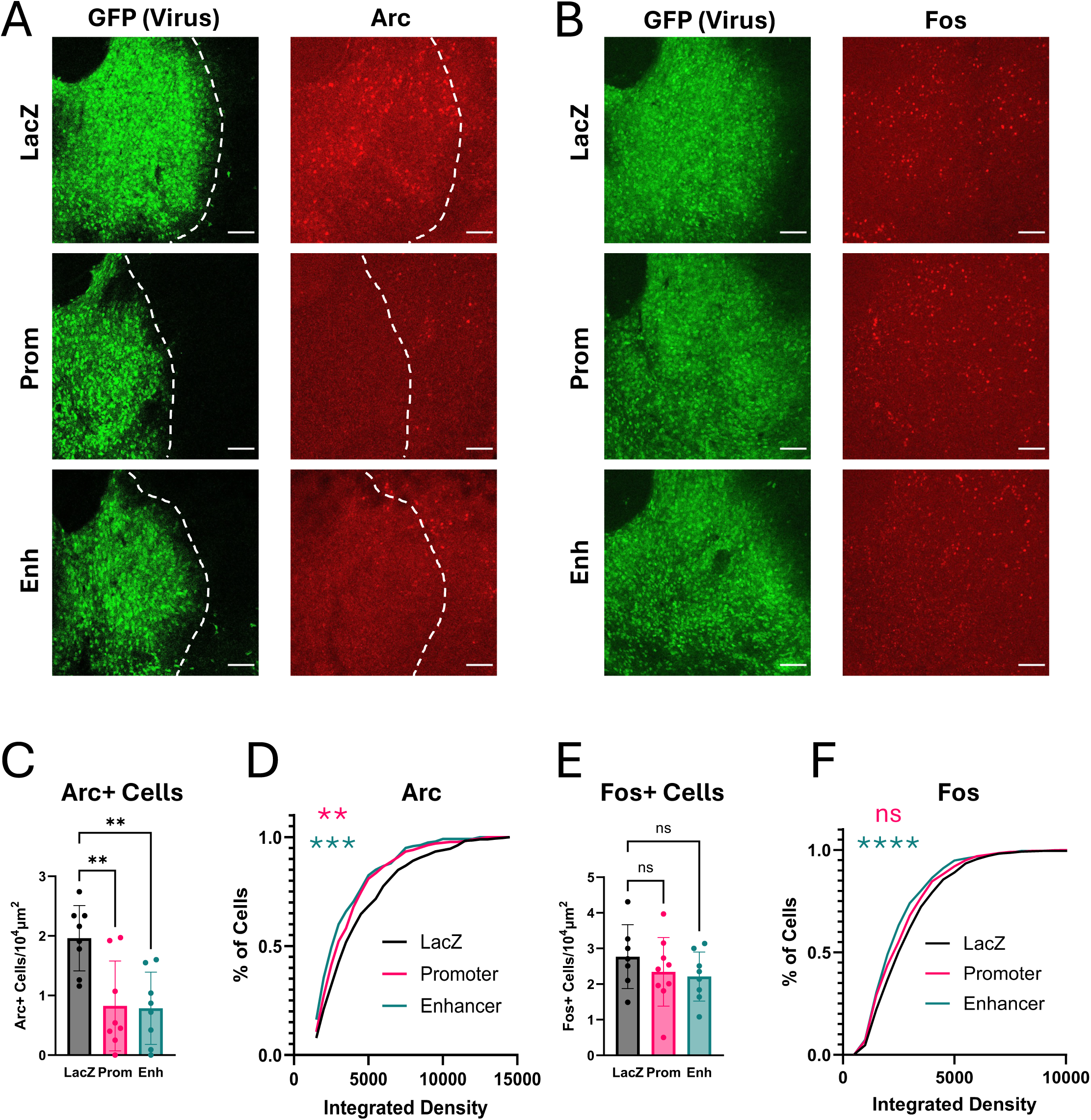
Repression of *Arc* enhancer, but not promoter, decreases Fos expression in Fos+ cells. **(A)** dCas9-KRAB mice from Figure 8 were injected with 20mg/kg cocaine, perfused one hour later, and brains were harvested for Arc and Fos staining on adjacent slices (red). Representative 20x NAc images showing Arc knockdown after repression of either the *Arc* promoter or the *Arc* enhancer. Scale bars are 100μm. **(B)** Same as (A) for Fos expression. **(C)** Average number of Arc+ cells in NAc per animal. **(D)** Cumulative distribution plot showing the integrated density changes of Arc+ cells between conditions. Cells are pooled by condition across animals. **(E)** Average number of Fos+ cells in NAc per animal. **(F)** Same as (D) for Fos expression. **p<0.01, ***p<0.001, ****p<0.0001.

When we quantified expression per cell, we found the expected decrease in the integrated density of the Arc+ cells with both regulatory element targets compared to the non-targeting LacZ control (**Figure10D**, *LacZ* – n = 470 cells, *Arc* Promoter – n = 272 cells, *Arc* Enhancer – n = 240 cells; Kolmogorov-Smirnov: *LacZ* vs Prom – p=0.0048, KS D = 0.1322; *LacZ* vs Enh p=0.0001, KS D = 0.1747, **Extended data Figure 4A, B**). Interestingly, however, there was also a significant decrease in the intensity of Fos immunostaining only in mice where the *Arc* enhancer was inhibited (**Figure 10F**, *LacZ* – n = 569 cells, *Arc* Promoter – n = 724 cells, *Arc* Enhancer – n = 582 cells; Kolmogorov-Smirnov: *LacZ* vs Prom – p=0.0565, KS D = 0.0748; *LacZ* vs Enh p=0.0001, KS D = 0.1463; **Extended data Figure 4C, D**). This is important because Fos is not just a marker of activated cells; it is a transcription factor, and the levels of Fos induction in a given cell impact the functional adaptations of that cell to stimuli (Chen et al., 2019; Gemberling et al., 2021). Thus, although repression of the *Arc* enhancer does not affect the number of Fos+ cells, it impacts the ensemble through decreases in the amount of Fos per cell. We discuss below the possible implications of this finding for future studies of Arc function.

## Discussion

In this study we developed and validated a set of CRISPR tools to control expression of the activity-inducible gene *Arc* in neurons. We confirmed that the *Arc* enhancer is selectively engaged for stimulus-induced expression of *Arc*, and we demonstrated that inhibiting either basal or activity-regulated *Arc* transcription is sufficient to decrease Arc protein expression in vivo. We used conditional CRISPR inhibition of Arc in visual cortex or NAc of adult dCas9-KRAB mice to test the requirements for Arc in plasticity of orientation tuning and cocaine-induced place preference, respectively. We anticipated that comparing the consequences of *Arc* regulation by its promoter or enhancer between these two brain regions might suggest common cellular consequences of Arc induction in the adult brain. However, despite evidence for the robust efficacy of our CRISPR inhibition strategy, we saw limited effects of Arc loss on our plasticity measures. These data show that stimulus-dependent induction of Arc expression does not always predict a requirement for Arc in experience-dependent plasticity.

### CRISPR confirms Arc enhancer mediates stimulus-selective regulation of Arc

Activity-responsive TFs bind to the *Arc* enhancer rather than its promoter, suggesting that the enhancer plays a central role in activity-inducible expression of *Arc* (Kawashima et al., 2009; Schaukowitch et al., 2014). Consistent with this model, we observed that targeted repression of the *Arc* enhancer led to decreases in both stimulus-induced eRNA and mRNA with no effect on *Arc* mRNA or protein levels at baseline. By contrast, recruitment of a CRISPR inhibitor or activator to the *Arc* promoter was sufficient to modulate Arc expression even in the absence of stimulation. In vivo, repression of either the enhancer or the promoter was sufficient to significantly reduce Arc protein expression in both V1 and the NAc, showing the substantial contribution of transcriptional regulation to Arc protein levels. However, in both regions, promoter-targeting gRNAs more strongly reduced Arc protein than enhancer targeting gRNAs, suggesting there may be a significant contribution of stimulus-independent expression to overall Arc levels in vivo. These data build on prior studies showing that enhancer targeting can be an effective way to assess the functional importance of neuronal activity-dependent gene regulation in the brain (Chen et al., 2019; Joo et al., 2016; Roethler et al., 2023).

### Limited effects of Arc knockdown on representational drift in adult V1

Neurons in sensory cortex must strike a balance between ongoing plasticity and stability of representations over time (Marks & Goard, 2021). Visual cortex presents a rich system for studying the functional consequences of synaptic plasticity on representational features including binocularity, orientation selectivity, direction selectivity, and receptive field location. Arc is widely expressed in adult visual cortex (**Figure 3**), where it undergoes daily light-dependent regulation of its expression (Kushinsky et al., 2024), and a prior study showed that spike timing-induced receptive field plasticity of visual cortical neurons was associated with Arc-dependent heterosynaptic plasticity (El-Boustani et al., 2018). We chose representational drift of orientation tuning over days as a model to test molecular mechanisms that balance plasticity with stability in the adult brain (Bauer et al., 2024). We predicted that heterosynaptic plasticity functions of Arc might be required to maintain tuning over time, moving AMPARs from weak to strong synapses to maintain orientation preference. We were able to observe substantial drift over a week (>10 degrees in preferred orientation) in a subset (∼25%) of V1 neurons validating our ability to detect ongoing plasticity of orientation preference. However, CRISPR inhibition of *Arc* neither reduced nor enhanced drift over a period of days.

Technical factors may have limited our outcomes in this study. First, in adult V1 the extent of orientation plasticity we detected is small, which is consistent with previous work (Bauer et al., 2024). If Arc is playing a relatively modest role regulating drift, our study may not be able to resolve the magnitude of this effect. Second, light induces Arc most highly in layers 4 and 6 of visual cortex (Jenks et al., 2017), whereas our calcium imaging is restricted to cells in layers 2/3. Layer 4 sends feedforward projections up to layer 2/3 (Callaway, 1998). Arc-dependent plasticity originating in layer 4 might have been missed without direct observation.

A prior study that performed calcium imaging in the binocular region of V1 similarly showed no effect on baseline orientation selectivity, though they did not assess drift (Jenks & Shepherd, 2020). Importantly, they did find that conditional loss of Arc in the adult brain impairs maintenance of binocularity. The differences in Arc sensitivity of these visual cortical features may arise from the way that these response properties are determined.

Both in development and in the adult brain, binocularity is regulated by Hebbian plasticity, in which strengthening of the relatively weak synapses from ipsilateral inputs to any given V1 neuron is tightly regulated by the activity of the stronger contralateral inputs to the same cell (Smith & Trachtenberg, 2010). By contrast, orientation selectivity in V1 neurons is more of a network process that arises not from the relative strength but the number of active synapses that share orientation selective firing properties (Jia et al., 2010; Scholl et al., 2021). Sensitivity to Arc might help to identify the types of cortical features that rely on Hebbian plasticity in the adult brain.

### Repression of the Arc promoter or enhancer in the NAc has minimal effects on CPP

Transcriptional regulation of synaptic plasticity has been widely studied as a mechanism for the behavioral adaptations induced by repeated exposure to the psychostimulants cocaine and amphetamine (Nestler, 2001). Arc is robustly induced in the NAc by cocaine (Salery et al., 2017) and reactivation of neurons labeled by ArcTRAP upon acute cocaine exposure was shown to enhance CPP (Salery et al., 2025). This shows that the population of neurons that induce Arc is important for behavioral responses to cocaine, but the requirement for cocaine-inducible Arc expression in CPP is less well understood.

Interpretation of altered CPP in the GFP knock-in strain of *Arc* KO mice (Penrod et al., 2020; Salery et al., 2017) is confounded by evidence for developmental wiring deficits in the dopamine system in this mouse strain (Gao et al., 2019; Managò et al., 2016). One study reported that shRNA-mediated KD of Arc in the NAc reduced CPP following a relatively low dose (5mg/kg) of cocaine, though significance was observed only in male mice, suggesting a relatively weak or context-specific effect (Wood et al., 2024).

We saw no significant effect on CPP when we repressed Arc induction in the NAc using gRNAs that targeted dCas9-KRAB to either the promoter or the enhancer of *Arc*. Notably, we used a higher dose of cocaine (20mg/kg) and more pairings of cocaine with the chamber compared with Wood et al (2024), so it is possible that we obscured a modulatory effect of Arc loss on the level of preference. Notably, Wood et al (2024) reported that shRNA KD of Arc in the adult NAc did not change the mini excitatory postsynaptic potentials or AMPAR currents again suggesting induced Arc may be unrelated to Hebbian plasticity in this context.

As an alternative way to test the function of NAc Arc in NAc-relevant plasticity, we conducted a novel object recognition memory task, assessing both short-term and long-term memory phases. Prior studies of hippocampal-mediated behaviors have indicated that adult expression of Arc is required for memory but not learning (Gao et al., 2018), and we expected our two-phase object recognition task might distinguish those stages.

However, we saw no effect of CRISPR-mediated Arc KD on object discrimination at either time point. These results again differ from the published effects of shRNA-mediated Arc KD in NAc where impairment of novel object recognition was observed, though the authors did not test recognition memory (Penrod et al., 2019). One limitation of our object recognition study is that it was performed after mice had been exposed to cocaine in CPP. Prior access to cocaine can alter recognition memory performance (Briand et al., 2008), though in general it would be expected to impair performance, not rescue a deficit.

### Arc function in Fos ensembles

Despite the absence of CPP or object recognition memory deficits in mice lacking Arc in the NAc, we did observe a reduction in the levels of Fos expression within the cocaine-induced NAc ensemble when we repressed the *Arc* enhancer. This is interesting because small ensembles of neurons activated by a stimulus like cocaine are thought to represent the experience for later recall (Josselyn et al., 2015). In the NAc, stimulus-induced Arc and Fos expression often overlap in the same sparsely-distributed neurons (Salery et al., 2025; Thibeault et al., 2025). We observed similar numbers of Fos expressing neurons when we inhibited Arc expression compared with control brains indicating that we have not disrupted the overall ensemble structure of this brain region. This may explain why CPP and object recognition memory were intact.

However Fos and Arc are more than just markers of ensembles - expression of these genes induces cellular adaptations that are likely to underlie memory retention (West, 2025). The changes in the level of Fos expression we observed, which were selective for mice in which we impaired the activity-inducible regulation of Arc by targeting the enhancer, raise the possibility that Arc may act to modify the function of ensemble neurons selectively in the context of a plasticity stimulus. Interestingly, in a recent study, Coda et al (2025) used a Fos-inducible dCas9/CRISPR strategy to regulate Arc expression only in Fos-expressing cells of the dentate gyrus (DG) in the adult mouse hippocampus (Coda et al., 2025).

Although this manipulation targeted a very small fraction (∼1%) of cells, it was sufficient to alter behavior in a contextual fear conditioning task. These data raise the possibility that Arc might have functions in plasticity that are specific to ensemble neurons. Whether these behavioral changes arose from Arc-dependent changes in AMPAR trafficking was not addressed in the Coda et al (2025) study, though other competitive mechanisms including transcription-dependent changes in intrinsic excitability have been suggested to underlie ensemble formation and function (Han et al., 2007). Notably, Arc shows substantial nuclear localization following stimulus-dependent induction where it has been proposed to mediate transcriptional regulation (Korb et al., 2013), though the importance of this mechanism for its function in the adult brain remains a topic for future study.

## Acknowledgments

This was supported by NIH grant R21EY032335 (AEW and LLG) and the Paul G. Allen Frontiers Group (AEW and LG). We would like to thank the Duke Behavioral and Neuroendocrine Core, the Light Microscopy and Core Facility, and the Duke Viral Vector Core facilities for their services. We also thank Mariah Hazlett, Krisna Sinha, and Nolyn Mjema for their help in this project.

---

**Extended Data Figure 1.**
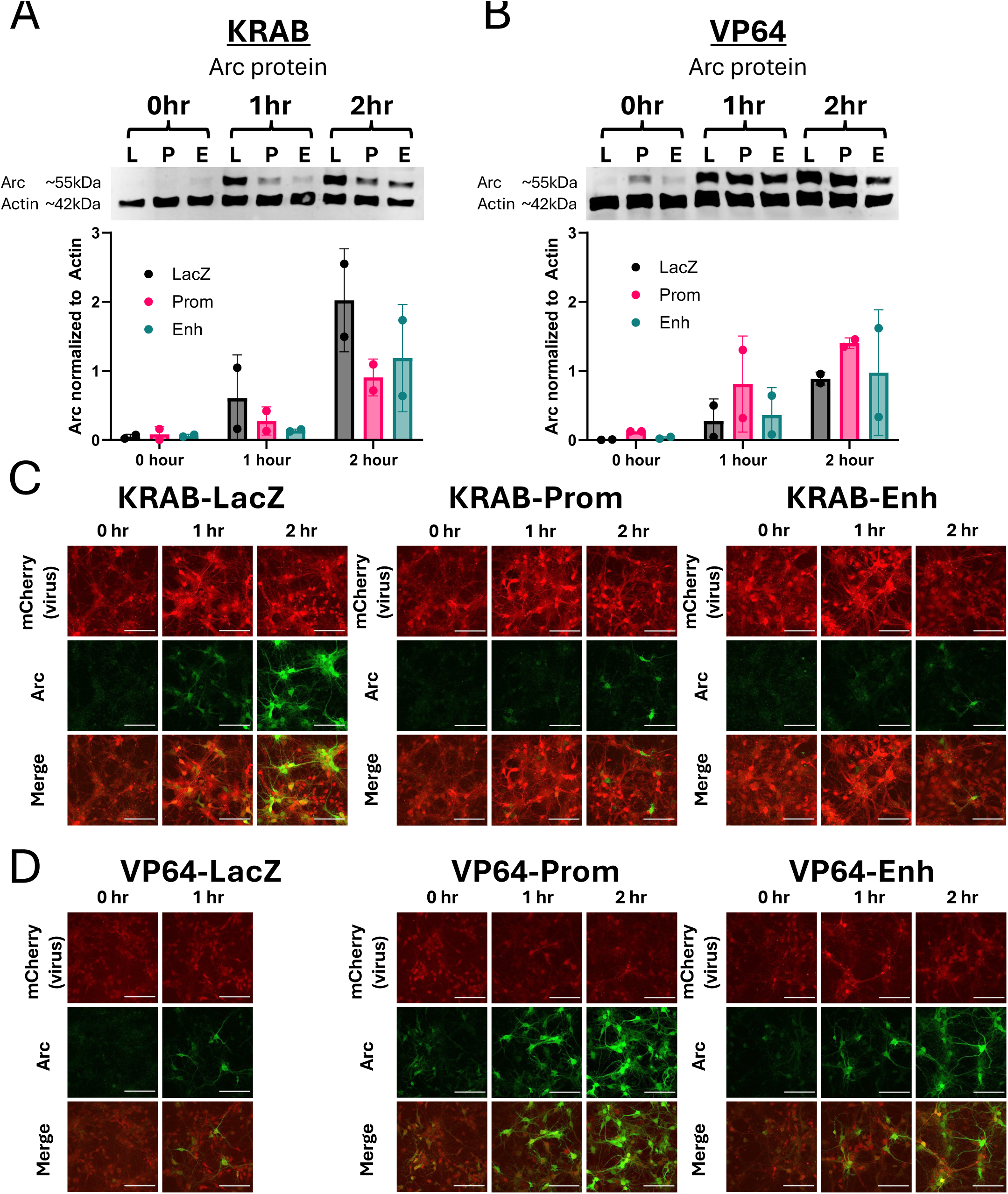
dCasG-KRAB and dCas9-VP64 modulate Arc protein expression in vitro. **(A, B)** Western blots from DIV7 cultured cortical neurons showing changes in Arc protein after targeting with dCas9-KRAB **(A)** or dCas9-VP64 **(B)** mCherry lentiviruses. Neurons were stimulated with BDNF for 0, 1, or 2 hours before harvesting. Data are represented as mean ± SD. No statistics were performed due to low sample size. **(C, D)** Immunocytochemistry images from cultured neurons on coverslips that were infected with dCas9-KRAB **(C)** or dCas9-VP64 **(D)** mCherry lentiviruses and stimulated with BDNF. Neurons were stained for Arc expression (green). Samples for the *LacZ* condition at the 2 hour timepoint were unrecoverable. Scale bars are 100μm.

**Extended Data Figure 2.**
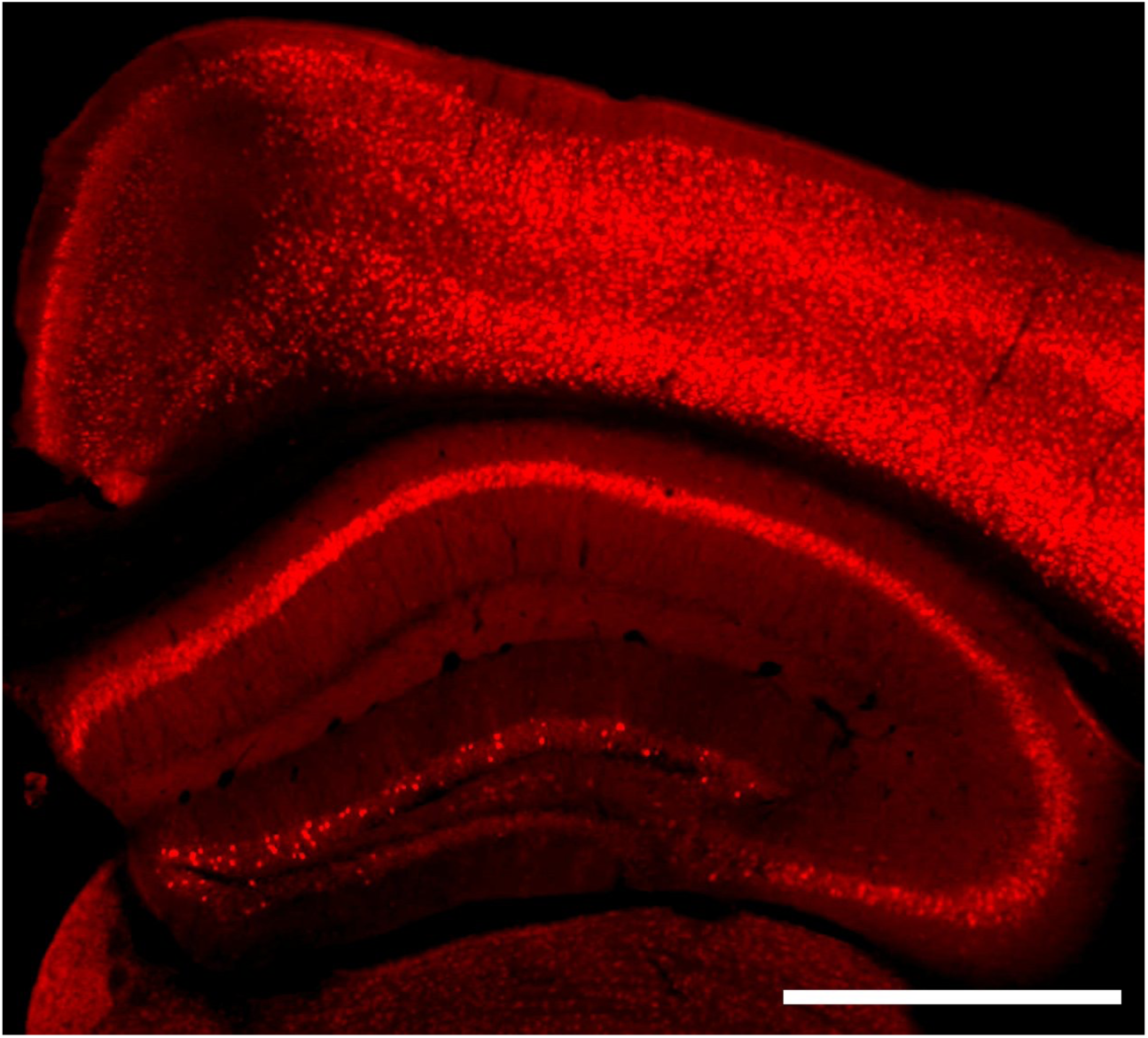
Arc expression varies in different brain regions. IHC image of mouse that underwent a CPP paradigm, a 20mg/kg cocaine challenge injection, and perfusion one hour later. Scale bar is 10cm.

**Extended Data Figure 3.**
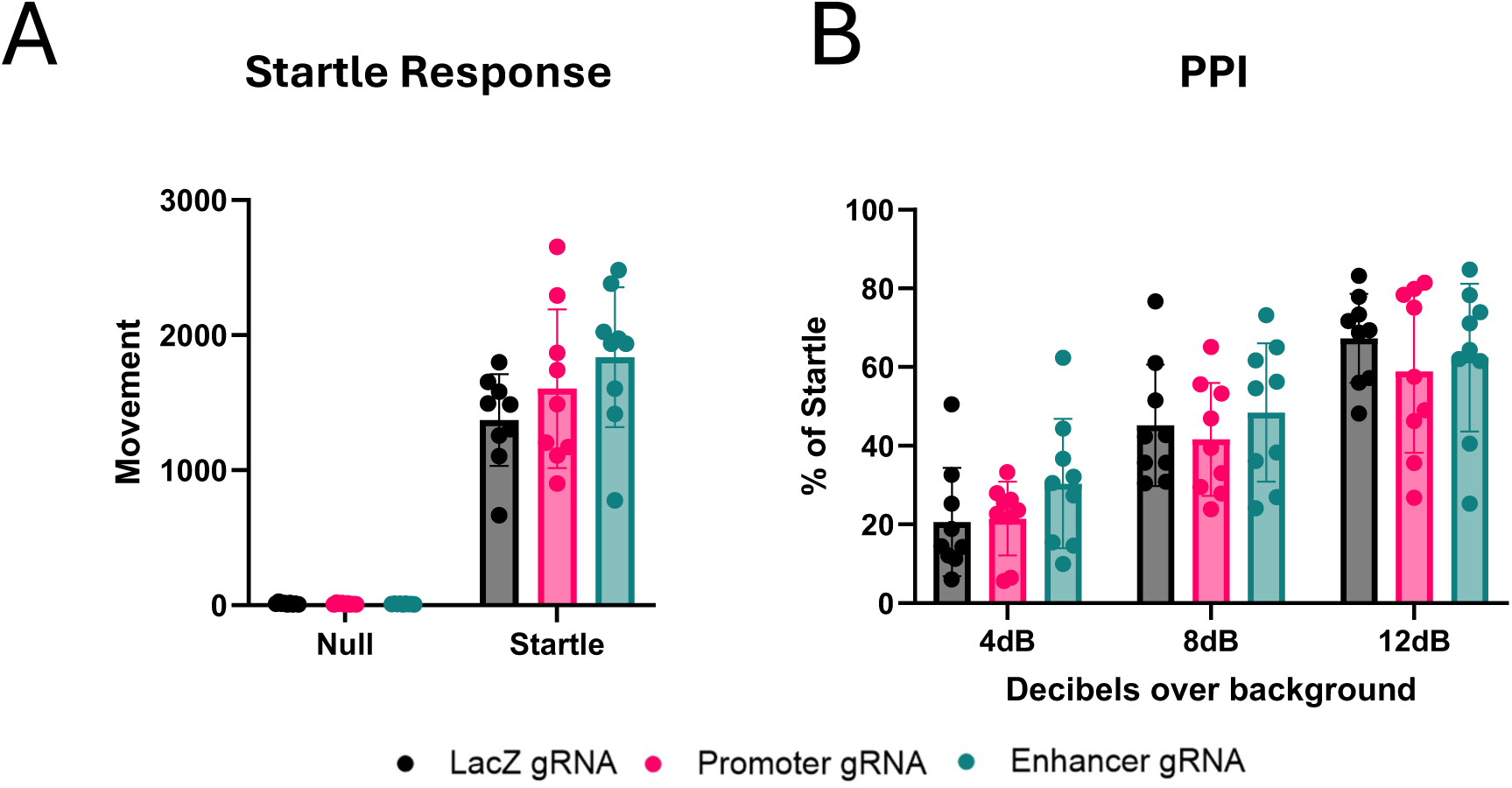
No changes in prepulse inhibition after Arc knockdown. **(A)** Startle responses to 120dB as baseline for PPI. Two-way rmANOVA, main effect of test: F(1,24) = 282.4, main effect of gRNA: F(2, 24) = 0.1607. **(B)** Amount of movement following pulse of sound, measured as percent of startle response. Two-way rmANOVA, main effect of dB: F(1.505, 36.11) = 91.84; main effect of gRNA: F(2, 24) = 0.5455. *p<0.01.

**Extended Data Figure 4.**
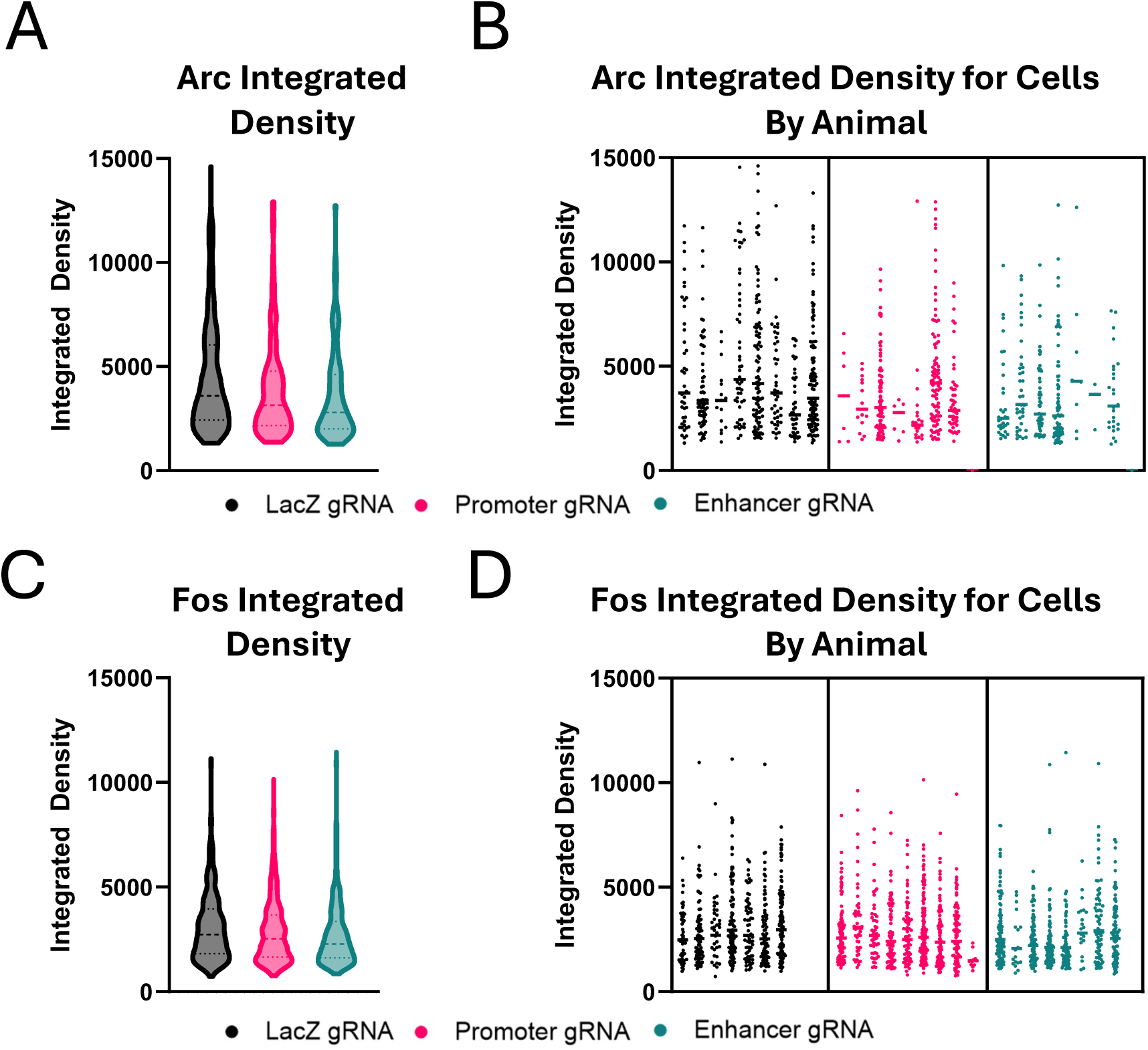
Expression levels of Arc and Fos ensembles. Mice were injected with 20mg/kg and perfused 1 hour later for immunostaining of Arc and Fos. **(A)** Violin plot showing the distribution of Arc expression from all Arc+ cells pooled across animals within each gRNA condition. **(B)** Nested distribution showing Arc expression of cells for each animal. **(C, D)** Same as (A, B) for Fos expression.

